# Global and genetic regulation of gene expression in human endothelial and vascular smooth muscle cells

**DOI:** 10.1101/2024.12.25.630318

**Authors:** Pengyuan Liu, Atrayee Ray, Yong Liu, Manoj K. Mishra, Qiongzi Qiu, Rajan Pandey, Bhavika Therani, Jing Huang, Jiayi Ren, Cary Stelloh, Xiaowen Bai, Andrew S. Greene, Allen W. Cowley, Aron M. Geurts, Sridhar Rao, Mingyu Liang

## Abstract

**Background:** The understanding of genetic and epigenetic regulation of gene expression in endothelial and vascular smooth muscle cells remains fragmented with limited experimental validation.

**Methods:** Chromatin conformation (Micro-C), DNA methylation (RRBS), chromatin accessibility (ATAC-seq), and transcriptome profiles (RNA-seq) were mapped in human induced pluripotent stem cell (hiPSC)-derived, isogenic endothelial and vascular smooth muscle cells (iECs and iVSMCs). CTCF and RAD21 were depleted to assess the functional relevance of chromatin architecture, and genome editing was used to evaluate the allelic effect of a blood pressure-associated single nucleotide polymorphism (SNP).

**Results:** Significant correlations were identified between gene expression levels and chromatin interactions, chromatin accessibility, and DNA methylation in iECs and iVSMCs, with chromatin interactions showing the strongest association. Chromatin contact regions displayed distinct epigenetic landscapes depending on the types of regulatory element interactions involved. Perturbation of CTCF and RAD21 revealed their differential regulatory effects, particularly on the expression of genes overlapping chromatin contacts, with RAD21 exhibiting a broader regulatory impact. SNPs associated with several vascular traits were enriched in chromatin loops or accessible regions in iECs or iVSMCs. Precise genome editing demonstrated allele-dependent effects of SNP rs9833313 on the expression of *SHOX2* located 247.4 kbp away but within the same chromatin loops as the SNP.

**Conclusion:** This study provides an extensive epigenetic landscape of vascular cells that may drive novel research on the role of genetic and epigenetic mechanisms of vascular function and disease as demonstrated by our targeted experiments.

## Introduction

Endothelial and vascular smooth muscle cells are the major cell types in blood vessels. Gene expression patterns in these cells underlie vascular function, and abnormal gene expression in these cells contribute to the development of a wide range of diseases that involve vascular abnormalities such as hypertension, coronary artery disease, and stroke.

Gene expression in endothelial and vascular smooth muscle cells, as in other cell types, may be regulated by epigenetic factors including DNA methylation, histone modifications, chromatin accessibility, and chromatin conformation. Previous studies such as the Encyclopedia of DNA Elements (ENCODE) project have generated some epigenomic profiles for vascular cells, mostly for human umbilical vein endothelial cells and aortic smooth muscle cells^1,2^. Insights gained from these epigenomic profiles have contributed to significant advances in our understanding of vascular biology and disease^3–5^. Genetic factors may also regulate gene expression. Thousands of single nucleotide polymorphisms (SNPs) have been associated with vascular traits and diseases. Most of these SNPs are in noncoding parts of the DNA ^6–8^. These noncoding SNPs may influence their associated traits by regulating gene expression in vascular cells through epigenetic mechanisms.

Several significant knowledge gaps remain in our understanding of the genetic and epigenetic regulation of gene expression in vascular cells. First, while some aspects of the epigenome in vascular cells, such as histone modifications, have been examined extensively, analyses of other aspects such as chromatin conformation and accessibility are limited. In addition, most of the available analyses were performed in cells with varying genomic backgrounds and in separate studies, preventing a rigorous, integrated analysis of epigenetic regulation.

Second, most epigenomic analyses have been correlative. The functional role of most epigenomic features in determining gene expression in vascular cells remains largely unproven. Direct manipulation of epigenetic regulators, followed by interrogation of global and nearby gene expression, is rare for vascular cells.

Third, despite efforts to map trait-associated SNPs to epigenetic features and associating SNPs with tissue-level gene expression using approaches such as expression quantitative trait locus (eQTL) mapping ^9,10^, the functional role for most noncoding SNPs in regulating gene expression remains unproven, especially in specific cell types. This is true for SNPs associated with all traits including vascular traits.

In this study, we utilized a multi-pronged, integrated approach to address these knowledge gaps. We began by mapping epigenomic landscapes in isogenic endothelial and vascular smooth muscle cells derived from one line of human induced pluripotent stem cells (hiPSCs). We integrated transcriptomes, DNA methylation, chromatin accessibility, and chromatin conformation to gain insight into the potential role of these epigenetic features and the coordination among them in gene regulation. We then knocked down CCCTC-binding factor (CTCF) and RAD21, which are key regulators of chromatin conformation, to identify genes sensitive to chromatin folding. Finally, we integrated the epigenomic data with genetic findings to nominate a noncoding SNP and used precise genome editing to identify the allelic effect of the SNP on gene expression and epigenetic features.

These findings provide novel insights into the global and genetic regulation of gene expression in human endothelial and vascular smooth muscle cells. Our approach of integrating epigenomic mapping with targeted intervention and precise genome editing can be applied to study any cell types and traits.

## Results

### Epigenomic profiles of hiPSC-derived endothelial and vascular smooth muscle cells

To unravel epigenetic and genetic control of gene expression in vascular cells, we constructed comprehensive epigenomic landscapes of hiPSC-derived endothelial cells (iEC) and vascular smooth muscle cells (iVSMC) (**Fig 1A**). This analysis integrates transcriptome profiling (RNA-seq), DNA methylation mapping (RRBS), chromatin accessibility analysis (ATAC-seq), and 3-D chromatin conformation mapping (Micro-C).

**Figure 1.**
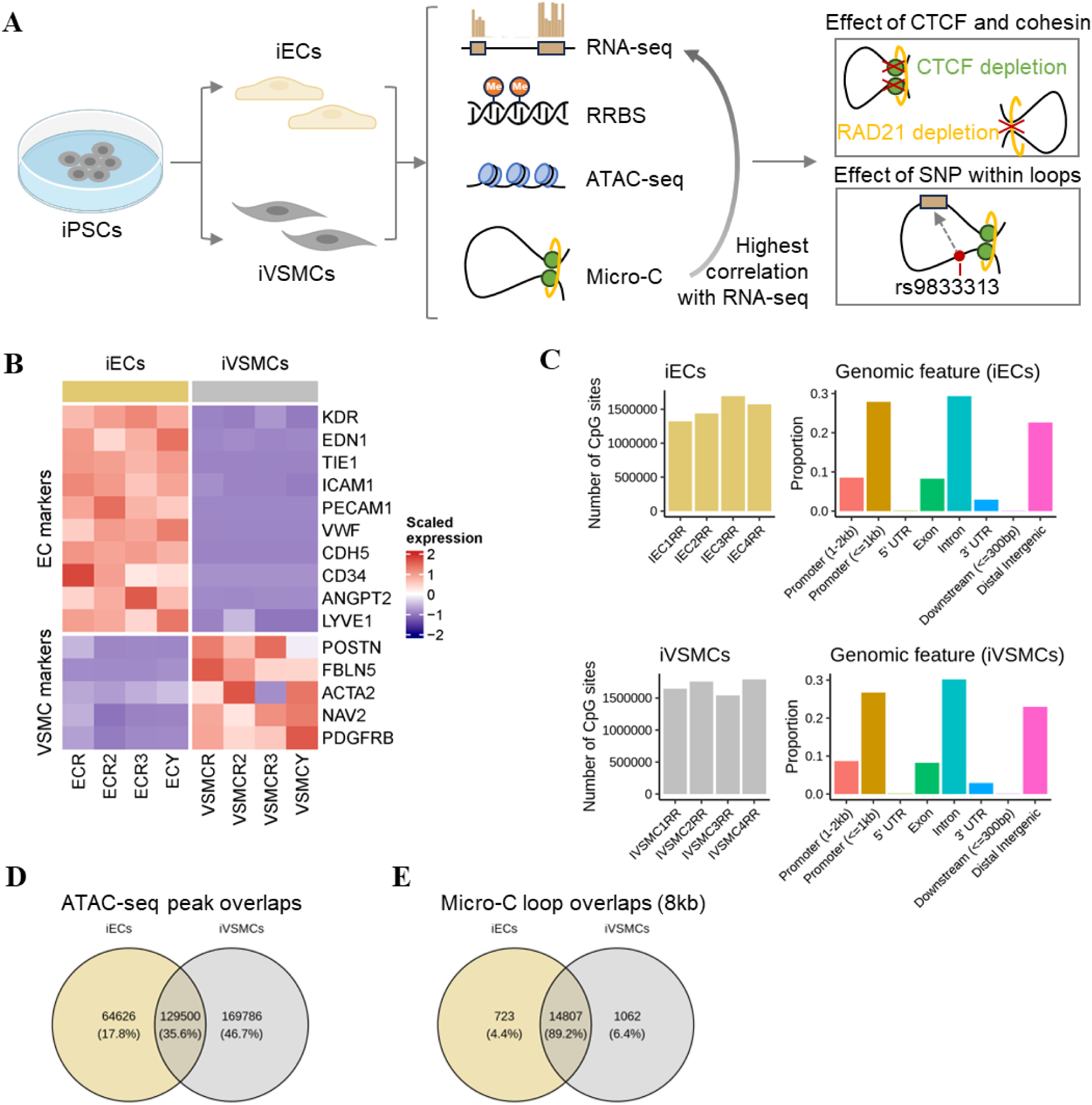
Overview of integrated analysis of epigenomic profiles in iECs and iVSMCs. (**A**) Study design. Human induced pluripotent stem cells (hiPSCs) were differentiated into isogenic endothelial and vascular smooth muscle cells (iECs and iVSMCs). Next, comprehensive epigenomic landscapes were constructed for iECs and iVSMCs, and transcriptomic, DNA methylation, chromatin accessibility, and chromatin conformation data were integrated to assess the role and coordination of these features in gene regulation. Then, chromatin conformation regulators CTCF and RAD21 were depleted to identify genes responsive to chromatin folding. Finally, integrated analysis of epigenomic and human genetic data was performed to investigate the impact of a noncoding SNP on gene expression and epigenetic features, followed by a proof of principle study using precise genome editing. (**B**) RNA-seq for profiling gene expression. Heatmap of selected marker genes of EC and VSMC is shown. Each column was an individual sample. (**C**) Reduced representation bisulfite sequencing (RRBS) for profiling genome-wide DNA methylation. Numbers of CpG sites detected at a coverage depth of ≥5X and their genomic distribution are shown. (**D**) Assay for Transposase-Accessible Chromatin with high-throughput sequencing (ATAC-seq) for profiling chromatin accessibility. Numbers of chromatin peaks detected in iECs and iVSMCs are shown. (**E**) Micro-C for profiling chromatin conformations. Numbers of chromatin loops detected at 8kb resolution in iECs and iVSMCs are shown.

Global gene expression profiling via RNA-seq revealed distinct expression patterns between iECs and iVSMCs, segregating the two cell types (**Fig S1A-B**, and **Table S1**). Differential expression analysis identified genes contributing to each cell type’s unique functions and identities (**Fig 1B**). Functional enrichment analysis of iEC-enriched genes highlighted biological processes associated with system development, particularly renal, urogenital, and limb development, as well as extracellular matrix organization (**Fig S1C** and **S1E**). Conversely, iVSMC-enriched genes were associated with Ras protein signaling and actin filament regulation. Pathway analysis further demonstrated that iEC-enriched genes were significantly involved in TGF-beta, Hippo, and Wnt signaling pathways, while iVSMC-enriched genes showed enrichment in MAPK, Rap1, and Ras signaling pathways (**Fig S1D** and **S1F**).

Genome-wide DNA methylation patterns were profiled in iECs and iVSMCs using Reduced Representation Bisulfite Sequencing (RRBS), which detected approximately 1.6 million CpG sites with at least 5X coverage in each cell type (**Fig 1C**, **S2A-B** and **Table S2**). The majority of these CpG sites were located in promoters, exons, introns, and distal regulatory regions (**Fig 1C**). The resulting data revealed distinct DNA methylation patterns between iECs and iVSMCs, capturing cell-type-specific methylation nuances (**Fig S2C-E**).

Chromatin accessibility was examined with ATAC-seq, identifying 194,126 and 299,286 accessible chromatin regions in iECs and iVSMCs, respectively (**Fig 1D** and **Table S3**). These accessible regions predominantly clustered around transcription start sites (TSS), with 36% overlapping between iECs and iVSMCs, and 18% and 47% unique to iECs and iVSMCs, respectively (**Fig 1D** and **S3A-C**).

Chromatin conformations were mapped using Micro-C, identifying 9,198, 15,869, and 12,978 chromatin loops in iECs and 7,359, 15,530, and 13,743 chromatin loops in iVSMCs at 4, 8, and 16 kb resolutions, respectively (**Fig 1E**, **S4A** and **Table S4**). Average loop sizes were 220, 459, and 797 kb in iECs and 246, 462, and 800 kb in iVSMCs for the 4, 8, and 16 kb resolutions, respectively (**Fig S4B**). The chromatin contacts forming these loops primarily spanned promoter regions and distal regulatory intergenic regions, underscoring their potential regulatory role (**Fig S5**).

These multi-faceted epigenomic profiles provide a detailed view of the regulatory landscape in iECs and iVSMCs, highlighting distinct epigenetic features of these isogenic cells.

### Distinct epigenomic features at representative genes essential for vascular function

Utilizing the epigenomic landscapes described above, we examined genes known to be essential for vascular function. We included *AGTR1* (Angiotensin II Receptor Type 1), *NOS3* (Nitric Oxide Synthase 3), and *ACE* (Angiotensin I Converting Enzyme) as representative examples. We tested the hypothesis that the epigenomic landscapes of these genes would reveal distinct regulatory features that likely contribute to their expression.

The expression of *AGTR1* was much more abundant in iVSMCs than in iECs, which was associated with elevated chromatin accessibility at the promoter region in iVSMCs, highlighted by prominent ATAC-seq peaks, indicating this region’s active regulatory status in iVSMCs (**Fig 2A** and **S6A**). By contrast, *NOS3* was much more highly expressed in iECs and displayed higher DNA methylation levels across the gene region in iVSMCs (**Fig 2B** and **S6B**). The *ACE* gene presented a loop exclusively in iEC and not in iVSMC, which connected its promoter to a super-enhancer identified from HUVEC data, a configuration likely facilitating enhanced gene expression in iECs by bringing distal regulatory elements into proximity with the promoter (**Fig 2C** and **S6C**).

**Figure 2.**
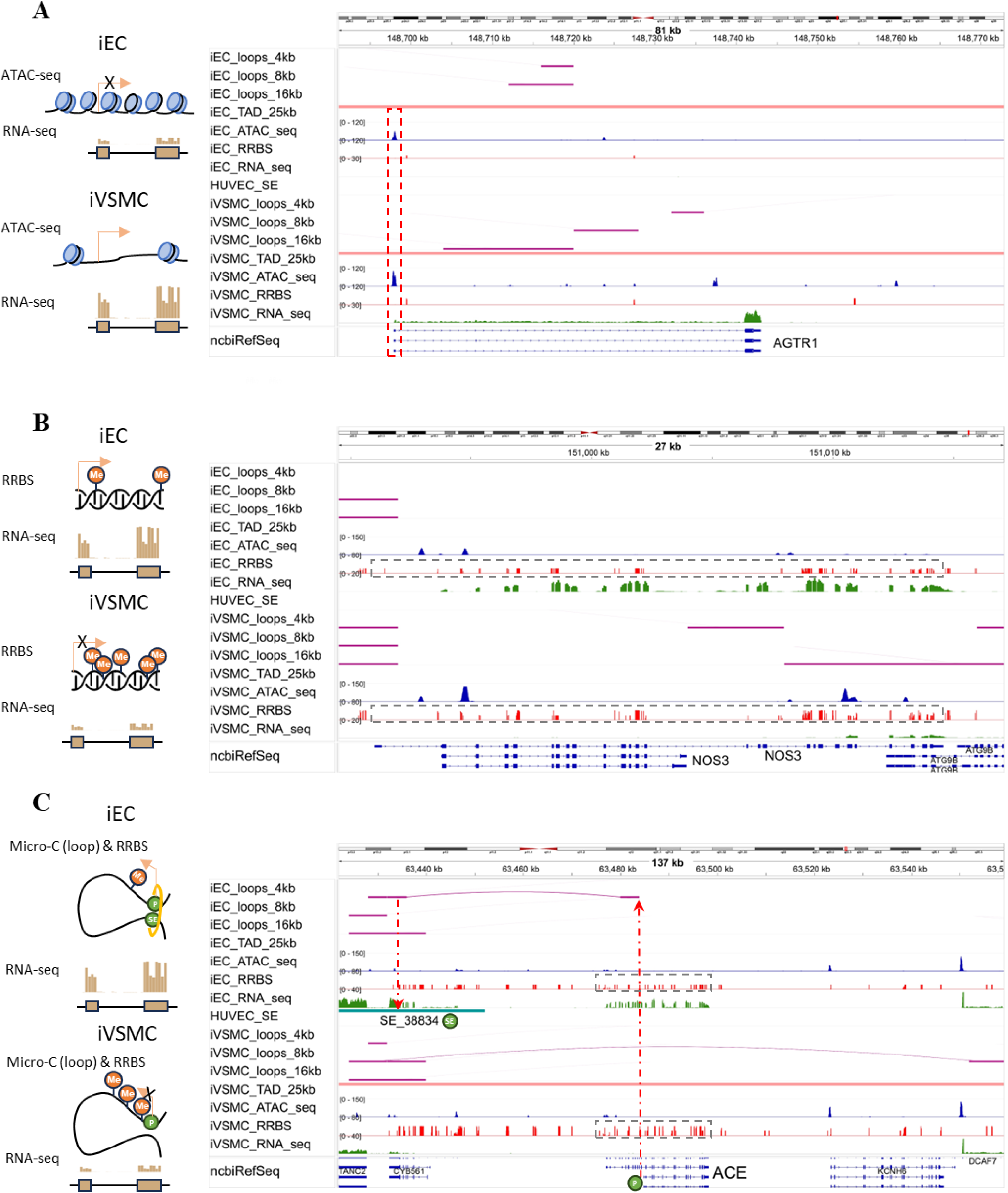
Epigenomic features in iECs and iVSMCs for representative genes important for vascular function. Genomic tracks show interaction loops and TADs (based on Micro-C), chromatin accessibility (ATAC-seq), DNA methylation (RRBS), and gene expression (RNA-seq) in iECs and iVSMCs, and super-enhancers obtained from dbSUPER. The schematic on the left depicts a proposed mechanism for the differential gene expression observed between iECs and iVSMCs. (**A**) AGTR1 (Angiotensin II Receptor Type 1). The vertical, red dashed rectangle highlights increased ATAC-seq peaks in the AGTR1 promoter region in iVSMCs. (**B**) NOS3 (Nitric Oxide Synthase 3). The horizonal, red dashed rectangle highlights elevated DNA methylation peaks in the NOS3 gene region in iVSMCs. (**C**) ACE (angiotensin I converting enzyme). Red arrows indicate the loop overlapping the ACE promoter and a super-enhancer identified in HUVECs. The horizontal, gray dashed rectangle highlights elevated DNA methylation levels in the ACE gene region in iVSMCs. See Supplemental Figure S6 for zoomed-out views of the AGTR1 and NOS3 regions, displaying TADs and loops, and zoomed-in views of the ACE gene region. TAD, topologically associating domain; HUVEC: Human umbilical vein endothelial cell; SE: super-enhancer. The Y-axis scale is identical across cell types to enable direct comparisons.

Together, these data illustrates how knowledge of distinct epigenetic features, such as chromatin accessibility, DNA methylation, and chromatin looping, may help to understand how the expression of such genes known to be important for vascular function is regulated.

### Global association of epigenetic features with gene expression

We expanded the analysis to genes in iECs and iVSMCs globally with a quantitative assessment of the associations between epigenetic features and gene expression levels.

Promoter DNA methylation is well-known to influence gene transcription by interfering with transcription factor binding, recruiting Methyl-CpG-binding domain proteins, and altering chromatin structure, typically leading to reduced gene repression ^11,12^. Consistent with this, we observed a statistically significant negative correlation between promoter DNA methylation levels and gene expression, though the correlation strength was modest (r=-0.06) (**Fig 3A**). In contrast, enhancers, essential distal *cis*-regulatory elements that work in tandem with promoters to regulate transcription, displayed a positive correlation between DNA methylation levels and nearby gene expression (**Fig 3B**). This data suggests distinct regulatory roles for methylation at promoters versus enhancers in gene expression in iECs and iVSMCs.

**Figure 3.**
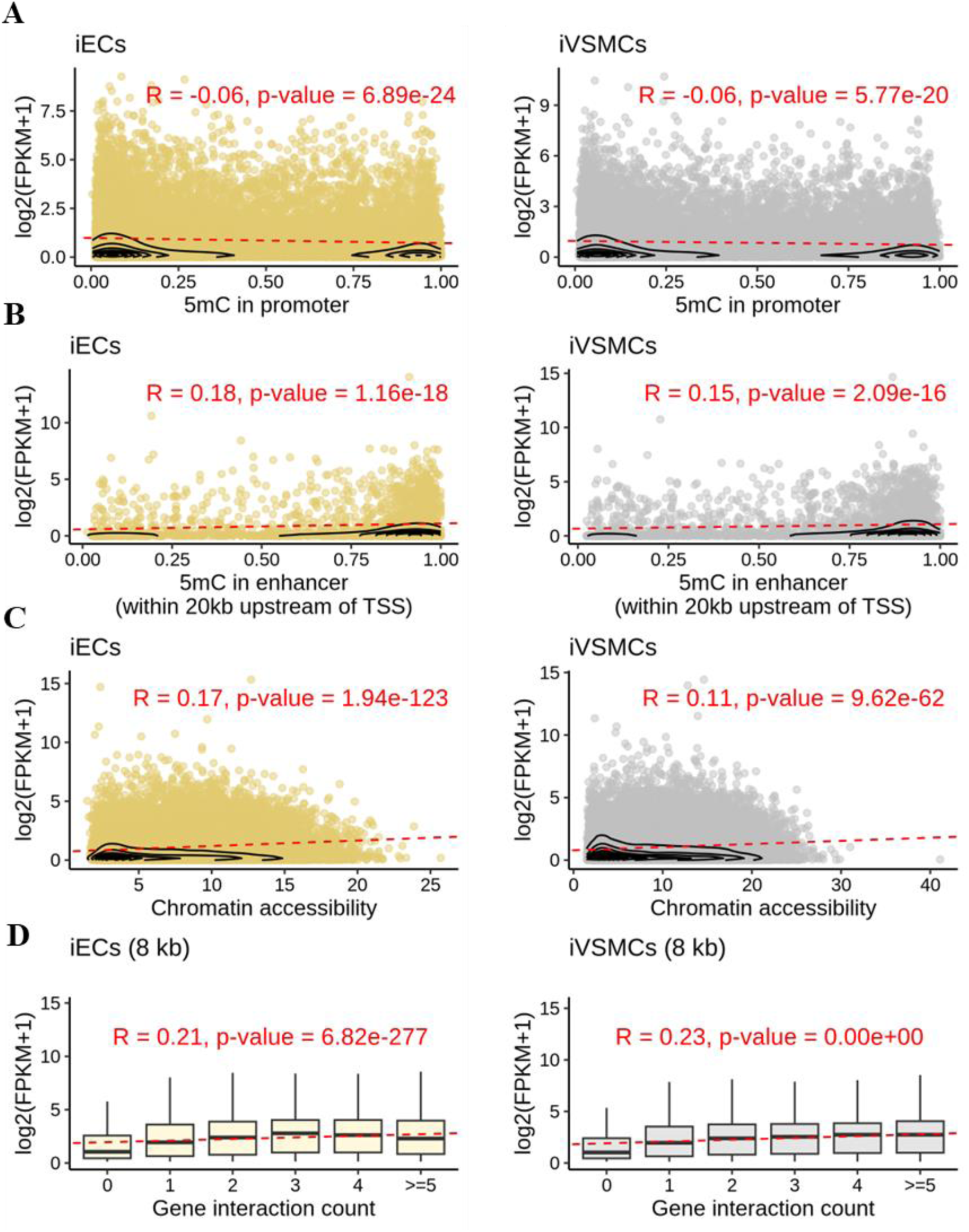
Epigenetic marks are associated with gene expression in iECs and iVSMCs. (**A, B**) Correlation between promoter/enhancer DNA methylation and nearby gene expression. Contour lines represent areas of similar density. (**C**) Correlation between chromatin accessibility and nearby gene expression. Contour lines represent areas of similar density. (**D**) Correlation between chromatin interaction counts and nearby gene expression. Spearman correlation was used to assess the relationship between epigenetic features and gene expression.

Chromatin accessibility across the genome reflects a complex network of physical interactions among enhancers, promoters, insulators, and chromatin-binding factors that cooperatively regulate gene expression ^13,14^. Our results indicate that increased chromatin accessibility is associated with higher expression levels of nearby genes (**Fig 3C** and **S7**).

Chromatin conformations play a regulatory role in gene transcription by bringing distal regulatory elements, such as enhancers, into proximity with promoters ^15^. Supporting this notion, we found that genes with a greater number of chromatin interactions within their gene bodies were more highly expressed in both iECs and iVSMCs (**Fig 3D**).

Overall, this integrated analysis of epigenetic and gene expression profiles indicate that epigenetic marks are significant predictors of gene expression levels in vascular cells.

### Regulatory elements in chromatin contact regions

In iECs and iVSMCs, the correlation of chromatin interaction counts with gene expression surpassed the correlation of DNA methylation or chromatin accessibility with nearby gene expression (**Fig 3**). Therefore, we focused subsequent analyses on further understanding the regulatory function of chromatin interactions in these cells. Chromatin interactions detected at 4 kb, 8 kb, and 16 kb resolutions showed comparable results (**Fig 4** and **S8-9**). For clarity, we primarily present our analyses based on the 8kb medium resolution.

**Figure 4.**
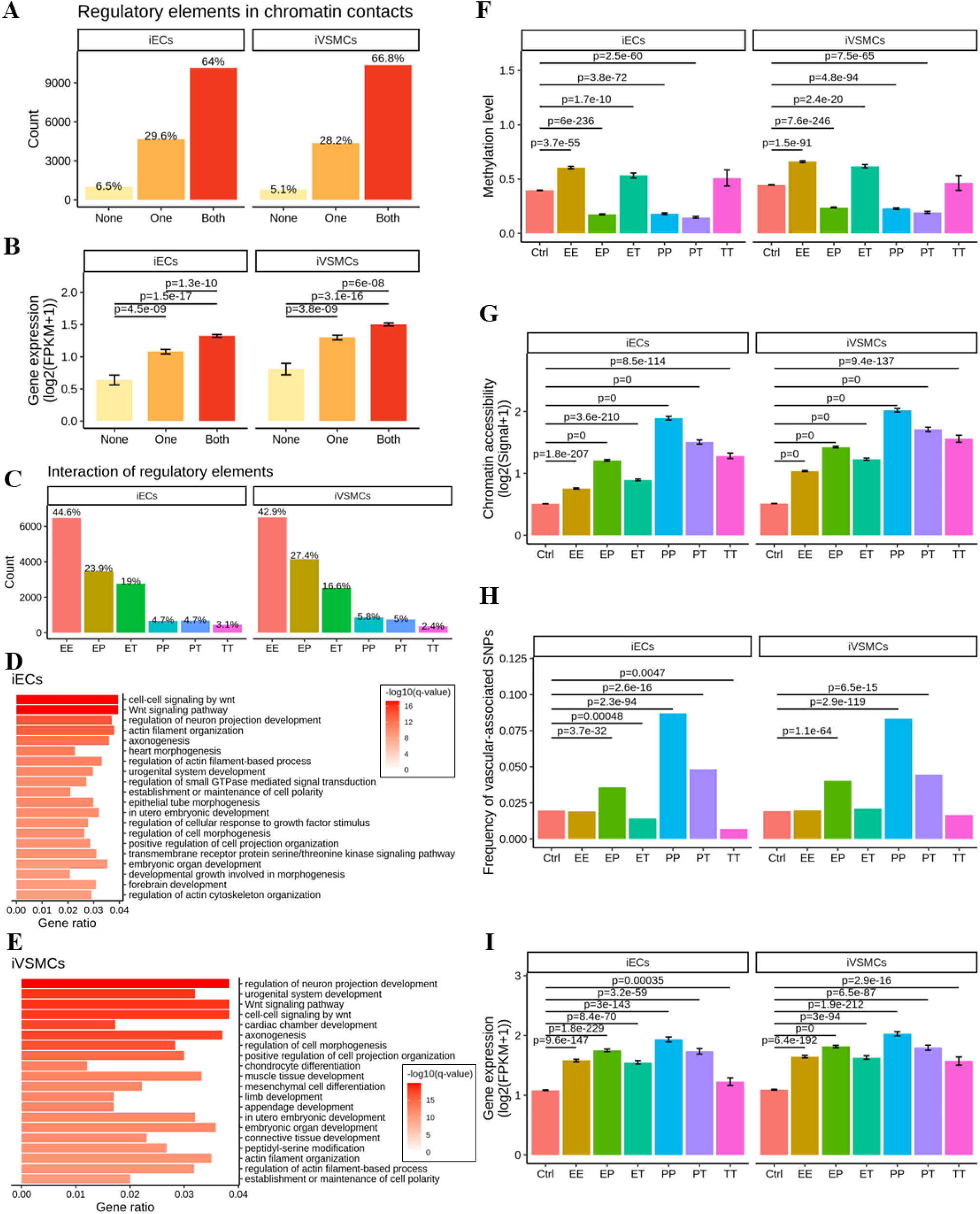
Chromatin contact regions in iECs and iVSMCs: regulatory element interactions, epigenetic features, and gene expression. (**A**) Chromatin contact regions of DNA loops were classified based on the presence of regulatory elements: “None” indicates that neither contact region of a loop contains a regulatory element; “One” indicates that one of the contact regions of a loop contains a regulatory element; and “Both” indicates that both contact regions of a loop contain regulatory elements. (**B**) Expression levels of genes located near chromatin contact regions with varying configurations of regulatory elements. (**C**) Distribution of interaction types between regulatory elements within chromatin contact regions. Key interaction types include promoter-promoter (PP), enhancer-promoter (EP), enhancer-enhancer (EE), enhancer-transcription factor binding site (ET), promoter-transcription factor binding site (PT), and transcription factor binding site-transcription factor binding site (TT) interactions. (**D, E**) GO and pathway enrichment analysis of genes near chromatin contact regions in iECs (**D**) and iVSMCs (E). (**F**) DNA methylation levels in chromatin contact regions with different types of regulatory interactions. (**G**) Chromatin accessibility in chromatin contact regions associated with various regulatory interactions. (**H**) Enrichment of vascular traits-associated SNPs in chromatin contact regions with different types of regulatory interactions. (**I**) Expression levels of genes near chromatin contact regions with distinct regulatory element interactions. Gene expression and epigenetic features are presented as mean ± standard error. Differences in quantitative variable between groups were assessed using Wilcoxon signed rank test, while differences in SNP proportions were evaluated using a two-sample proportion test.

We examined the types of known regulatory elements, as defined by Ensembl Regulatory Build (hg38), located within chromatin contact regions forming chromatin loops in iECs and iVSMCs. At 8kb resolution, more than 70% of the chromatin contact regions in iECs and iVSMCs overlapped with enhancers, 16% to 19% overlapped with promoters, and approximately 10% overlapped with transcriptional factor (TF) binding sites (**Fig S10**). Nearly 30% of chromatin loops in iECs and iVSMCs featured at least one regulatory element — enhancers, promoters, or TF binding sites — in one of the contact regions, while 64.9% of iEC loops and 66.8% of iVSMC loops contained regulatory elements in both contact regions (**Fig 4A**). Furthermore, genes near chromatin contacts containing regulatory elements exhibited significantly higher expression levels than genes near contacts lacking such elements (**Fig 4B**).

Chromatin contact regions exhibited various combinations of regulatory element interactions. In iECs, 44.6% of these were enhancer-enhancer (EE) interactions, 4.7% were promoter-promoter (PP) interactions, 3.1% were TF binding site-TF binding site (TT) interactions, 23.9% were enhancer-promoter (EP) interactions, 19.0% were enhancer-TF binding site (ET) interactions, and 4.7% were promoter-TF binding site (PT) interactions (**Fig 4C**). The distribution in iVSMCs was similar (**Fig 4C**). Additionally, approximately 80.9% of iECs and 81.7% of iVSMCs contact regions featured at least two types of interactions.

We further examined functional features of genes located within 5kb upstream and downstream of chromatin contact regions. In iECs, these genes are enriched in pathways related to development and morphogenesis, particularly Wnt signaling, axonogenesis, heart morphogenesis, and actin filament organization (**Fig 4D**). These findings suggest genes near chromatin contacts in iECs play a key role in regulating embryonic development and tissue morphogenesis. In iVSMCs, enriched pathways include Wnt signaling, neuronal projection development, cardiac and muscle tissue development, and actin filament organization, highlighting their involvement in vascular development and mesenchymal differentiation (**Fig 4E**). Both cell types share enrichment in pathways related to cellular polarity and growth factor responses, indicating their roles in regulating cell morphology and tissue structure.

Together, these data reveal the diversity and complexity of regulatory elements within chromatin contacts and their potential role in mediating the effect of chromatin interactions on gene expression in vascular cells.

### Epigenetic features of chromatin contacts involving regulatory element interactions

We further analyzed the epigenetic characteristics of chromatin contact regions involved in various regulatory element interactions. In both iECs and iVSMCs, chromatin contact regions involving PP, EP, and PT interactions exhibited lower DNA methylation levels compared to control regions (i.e., regulatory elements not located in chromatin contacts). Conversely, higher methylation levels were observed in chromatin contact regions involving EE, TT, and ET interactions (**Fig 4F**).

In general, chromatin contact regions involving regulatory element interactions demonstrated greater chromatin accessibility compared to control regions, with the highest accessibility observed in PP regions (**Fig 4G**). Additionally, SNPs associated with vascular traits were significantly enriched in chromatin contact regions involving PP, EP, and PT interactions, suggesting that these interactions are particularly relevant to the regulatory mechanisms influenced by genetic variation (**Fig 4H**). Genes located near chromatin contact regions involving PP, EP, and PT interactions also showed higher expression levels than those proximal to EE or other regulatory element interactions (**Fig 4I**).

These findings highlight how different regulatory element interactions within chromatin contact regions may function in coordination with other epigenetic features to influence gene expression.

### Role of nuclear architecture proteins in regulating iEC gene expression

Next, we experimentally tested the role of chromatin conformation in gene expression in iECs by targeting CTCF and the cohesin complex. CTCF and the cohesin complex, compromised of core subunits RAD21, SMC3, SMC1A, and STAG2, are critical mediators of nuclear architecture by forming and/or stabilizing chromatin loops. We hypothesized that depleting CTCF and the core cohesin subunit RAD21 would reveal their shared and unique contribution to chromatin-mediated gene regulation.

To test this hypothesis, we depleted CTCF and RAD21 in iECs using two lentiviral mediated shRNAs targeting each factor, resulting in an approximately 50% depletion of each protein **(Fig 5A**), although a greater depletion of mRNA was observed **(Fig S11A**). Despite similar knockdown efficiencies, RAD21 knockdown affected significantly more genes than CTCF knockdown based on RNA-seq (**Fig 5B** and **Table S5**). Approximately 30% of the genes affected by CTCF knockdown were affected by RAD21 knockdown, and 2.7% of the genes affected by RAD21 knockdown were affected by CTCF knockdown, indicating that these factors regulate largely distinct sets of genes in iECs (**Fig 5B**). To further understand how CTCF and RAD21 influence gene expression, we performed CUT&Tag experiments to identify the binding sites of CTCF and RAD21 in iECs. The analysis revealed that RAD21 occupied a greater number of genomic loci compared to CTCF, with RAD21-bound sites encompassing most CTCF-occupied sites but also additional sites, likely enhancers (**Fig 5C**). This finding provides a potential explanation for the broader regulatory effects observed upon RAD21 depletion, as its genome occupancy may directly influence more genes.

**Figure 5.**
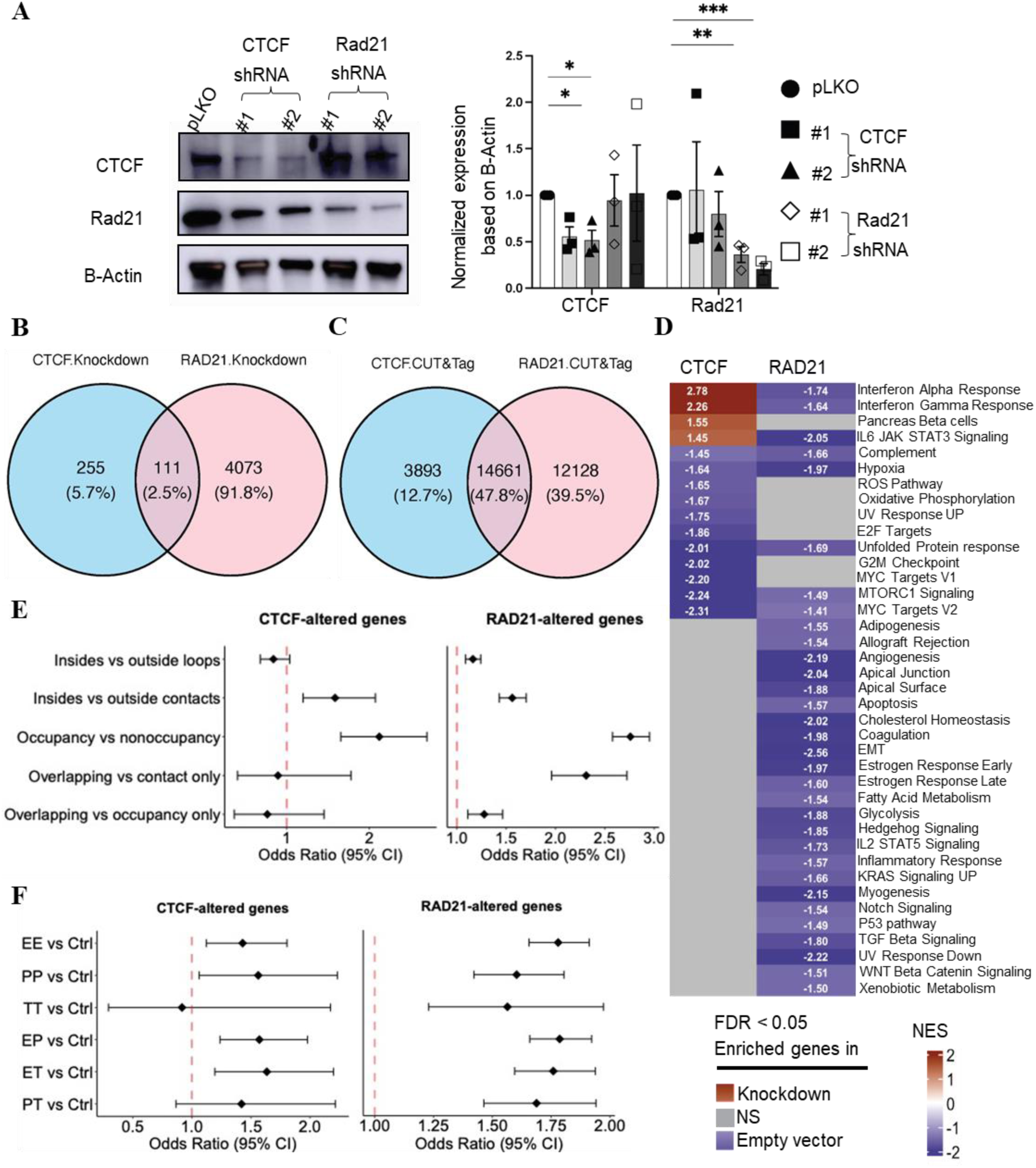
CTCF and RAD21 regulate gene expression in a chromatin organization-dependent manner. (**A**) Knockdown efficiency of CTCF and RAD21 in iECs, as determined by western blot. N=3 per shRNA; *, p<0.05, two-sided t-tests were used to assess expression difference between groups. (**B**) Number of differentially expressed genes (DEGs) in ECs following CTCF or RAD21 knockdown (**C**) Number of CTCF and RAD21 genomic occupancy sites in ECs, as detected by CUT&Tag. (**D**) Enriched pathways in CTCF/Rad21 depleted iEC compared to empty-vector-infected iEC control, identified by GSEA. All gene sets selected had an FDR<0.05. (**E**) Genes located in regulatory regions (such as DNA loops, chromatin contact, or near RAD21 or CTCF occupancy sites) are more likely to be altered in iECs, particularly following RAD21 knockdown. Overlapping represent overlapped regions between chromatin contacts and occupancy binding sites of CTCF or RAD21. (**F**) Genes near chromatin contacts involving regulatory interactions are more likely to be altered in response to CTCF or RAD21 knockdown in iECs.

Gene Set Enrichment Analysis (GSEA) of CTCF- and RAD21-depleted iECs revealed that the depletions altered distinct gene sets (**Fig 5D**). RAD21 depletion caused the downregulation of gene sets involved in the regulation of vascular function, including pathways related to TGFβ, Estrogen, and a broad array of inflammatory cascades such as Interferon α and γ, IL-6, IL-2, as well as the general gene set “Inflammatory Response”. In addition, EC-specific programs governing apical-basal cell polarity, coagulation, complement activation, and angiogenesis were also affected. In contrast, CTCF depletion induced the upregulation of gene sets associated with Interferon α and γ, and IL-6, reflecting an opposite effect compared to RAD21 depletion. Few pathways were shared between the two knockdowns, further supporting the notion that CTCF and RAD21 regulate distinct aspects of the iEC transcriptome. Consistent with this, gene ontology (GO) analysis of DEGs also revealed distinct roles for RAD21 and CTCF in gene regulation (**Fig S11B**). RAD21 knockdown was associated with morphogenesis and cytoskeletal organization, while CTCF knockdown was linked to neurodevelopment and metabolic regulation.

We examined the genomic features of the genes dysregulated in iECs following CTCF or RAD21 depletion. Given the role of CTCF and RAD21 in mediating chromatin contacts, we hypothesized that the dysregulated genes would be enriched in chromatin contact regions they occupied. Indeed, CTCF-a nd RAD21-sensitive genes in iECs were significantly more likely to overlap with chromatin contacts in iECs (**Fig 5E** and **S12A-B**). Moreover, RAD21-sensitive genes were more likely to be present within chromatin loops. As expected, genes near RAD21 or CTCF occupied sites were more likely to be altered upon depletion of these factors (**Fig 5E** and **S12A-B**), confirming direct effects of RAD21 and CTCF on the expression of genes in proximity to their binding sites. Furthermore, genes near chromatin contacts overlapping RAD21 occupancy were even more likely to be altered upon RAD21 knockdown, compared to those near contacts without RAD21 occupancy or those near RAD21 occupancy but not chromatin contacts, emphasizing RAD21’s role in contact-proximal regulation (**Fig 5E** and **S12A-B**).

To determine whether CTCF or RAD21 preferentially influence genes near chromatin contacts involving regulatory interactions, we compared genes near chromatin contacts with and without such interactions. Notably, genes near chromatin contacts involving regulatory interactions were more likely to be altered in response to RAD21 knockdown and, in most cases, CTCF knockdown, in iECs (**Fig 5F** and **S12C-D**), further highlighting the importance of chromatin context in the regulation of gene expression by these nuclear architecture proteins.

### Enrichment and functional impact of vascular traits-associated SNPs in epigenetic features in iECs and iVSMCs

Thousands of SNPs, including many in noncoding genomic regions, have been associated with vascular functions and diseases ^6,8,16,17^. We investigated whether SNPs associated with vascular traits are enriched in epigenetic features detected in iECs and iVSMCs and the potential connection between these SNPs and the expression of nearby genes. We found that numerous vascular traits-associated SNPs were located near expressed genes (RNA-seq), in more accessible chromatin regions (ATAC-seq), and/or within chromatin loops (Micro-C) in iECs and iVSMCs (**Fig 6A**).

**Figure 6.**
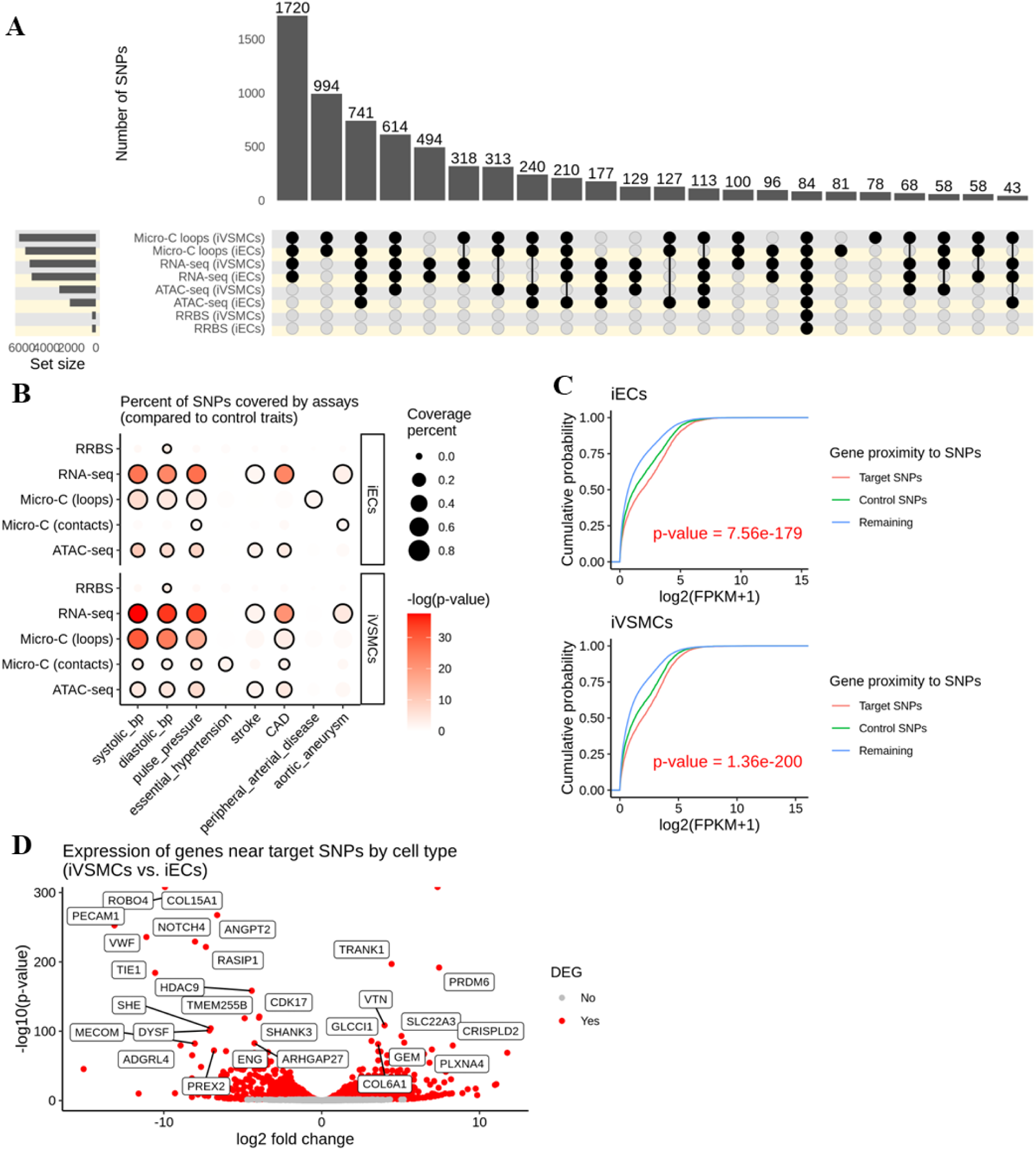
SNPs associated with vascular traits are enriched in epigenetic features in iECs and iVSMCs. (**A**) Upset plot showing the overlap of vascular traits-associated SNPs with epigenetic features identified by different omics assays. Black circles represent SNPs presented in the respective assays, while gray circles indicate their absence. (**B**) Significant enrichment of vascular traits-associated SNPs within regions of expressed genes (RNA-seq), chromatin-accessible areas (ATAC-seq), methylated promoters (RRBS), and DNA loops or contacts (Micro-C), compared with SNPs associated with non-vascular traits. Statistical significance was assessed using hypergeometric tests. (**C**) Elevated expression levels of genes located near vascular-associated SNPs. The difference in the probability density of gene expression across groups was analyzed using Kolmogorov-Smirnov tests. (**D**) Genes near vascular-associated SNPs exhibit cell-specific expression patterns in iECs and iVSMCs.

Remarkably, compared to SNPs associated with traits not directly or primarily related to blood vessels, SNPs associated with vascular traits were significantly more likely to be in epigenetic features that we detected in iECs and iVSMCs (**Fig 6B**). For example, SNPs associated with blood pressure (BP) phenotypes were more likely to be near expressed genes and in chromatin loops and accessible chromatin regions in both iECs and iVSMCs. SNPs associated with coronary artery disease (CAD) were more likely to be near expressed genes and in accessible chromatin regions in iECs and iVSMCs as well as in chromatin loops and contacts in iVSMCs. The specific enrichment in iVSMCs aligns with the important role of VSMCs in CAD, including their involvement in the buildup of atherosclerotic plaques, a hallmark of CAD. SNPs associated with stroke were more likely to be near expressed genes and in accessible chromatin regions in iECs and iVSMCs. SNPs associated with peripheral artery disease (PAD) are more likely to be in chromatin loops in iECs, aligning with the central role of ECs in mediating blood flow reduction characteristic of PAD (**Fig 6B**). Genes near vascular traits-associated SNPs showed significantly higher expression levels compared to those near SNPs associated with non-vascular traits or not near any of these SNPs, in both iECs and iVSMCs (**Fig 6C**). Additionally, a total of 1,489 genes (42%) near vascular traits-associated SNPs showed cell-type-specific expression patterns between iECs and iVSMCs (**Fig 6D**).

These findings suggest that vascular-associated SNPs are frequently located in chromatin loops and accessible chromatin regions in vascular cells, suggesting these SNPs may influence gene expression in vascular cells through epigenetic mechanisms, potentially linking them to vascular phenotypes in a cell-type-specific manner.

### BP-associated SNP rs9833313 influences the expression of SHOX2 247.4 kbp away in human vascular cells

As proof of principle, we performed experiments to test the effect of one of these vascular traits-associated SNPs located in epigenetic features of vascular cells. Our initial analysis identified 186 BP-associated SNPs enriched in Micro-C loops and accessible chromatin regions in both iECs and iVSMCs (**Fig 6B**). For the ease of genome editing, we narrowed these SNPs down to 33 that were not in linkage disequilibrium with any other SNP based on TOP-LD ^18^. Further selection reduced this list to 10 that were located at least 100 kbp from any protein-coding gene as such SNPs likely influenced their effector genes through epigenetic mechanism but their effector genes were difficult to identify (**Fig S13A**). Of these, 9 shared a chromatin loop with protein-coding genes in both iEC and iVSMC, and 3 were associated with more than one BP trait: rs9833313 and rs11688682 with systolic BP and pulse pressure, and rs76059372 with systolic and diastolic BP.

We performed the proof of principle study on rs9833313. The A allele of rs9833313, compared with the T allele, is associated with lower systolic blood pressure and pulse pressure in European population ^19^ and has an allele frequency of approximately 24% in Europeans. This SNP, located at Chr3:157859002 (hg38), resides 247.4 kbp from the transcriptional start site (TSS) of *SHOX2* (Short Stature Homeobox 2) and 236.9 kbp from its 3’ end (**Fig 7A).** Other neighboring genes include *SLC66A1L*, *VEPH1*, *PTX3*, *RSRC1*, and *MLF1*, with rs9833313 residing within in the same chromatin loops and TAD as *SHOX2*, *RSRC1*, and possibly other genes in both iECs and iVSMCs (**Fig 7B**).

**Figure 7.**
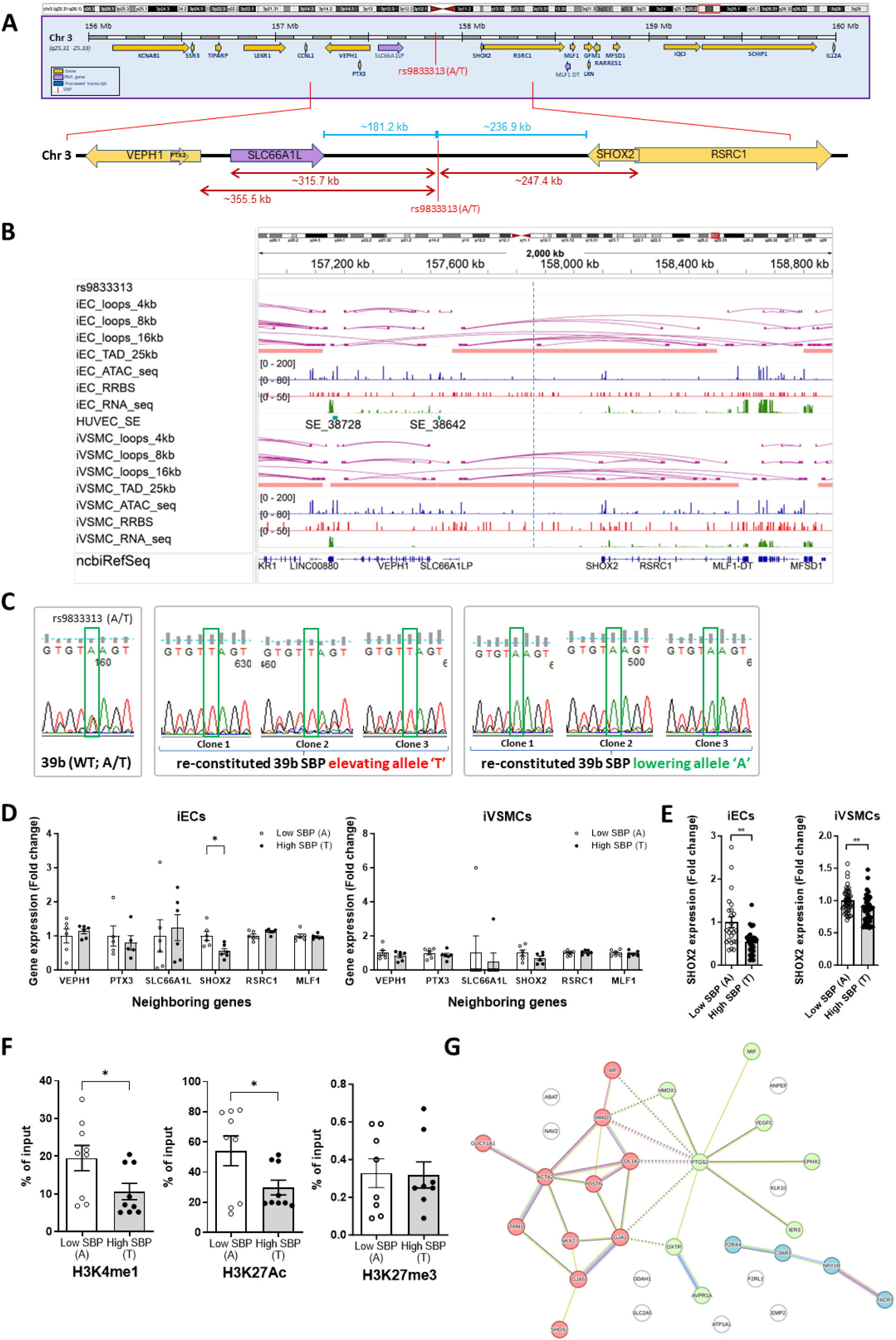
Allelic effects of BP-associated SNP rs9833313 on the expression of genes hundreds of kbp away. (**A**) Genomic context of SNP rs9833313, showing its location and surrounding region. (**B**) Chromatin loops and TADs (based on Micro-C) and other epigenetic features around rs9833313 in iECs and iVSMCs. The vertical dash line marks the SNP location. (**C**) Sanger sequencing confirmation of hiPSC lines edited to contain homozygous T (high-SBP) or A (low-SBP) allele at rs9833313. (**D**) Allelic effects of rs9833313 on the expression of local genes in iECs and iVSMCs, based on RNA-seq analysis. N=6 (3 clones of hiPSCs of each genotype, differentiated in 2 separate rounds), *, p<0.05. (**E**) Allelic effects of rs9833313 on the expression of *SHOX2* located 247.4 kbp away in iECs and iVSMCs, based on qPCR analysis. N=24-25 for iECs and n=45 for iVSMCs (3 clones of each genotype, differentiated in multiple separate rounds), **, p<0.01, two-sided unpaired t-test. (**F**) ChIP-qPCR analysis of histone mark binding at the rs9833313 region in iECs. N=9 for H3K4me1 and H3K27Ac and 8 for H3K27me3, *, p<0.05, two-sided unpaired t-test. Bars represent mean ± SEM. (**G**) STRING analysis showing association of SHOX2 and blood pressure physiology genes.

We specifically and precisely edited the allelic sequence of rs9833313 in hiPSCs. This was achieved by first deleting a 185 bp segment around the SNP and then reconstituting the SNP with the desired sequence, generating two isogenic hiPSC lines containing homozygous A (low SBP) or T (high SBP) allele at the rs9833313 site (**Fig 7C** and **S13B-E**). Three clones of hiPSCs were established for each line.

We differentiated the edited hiPSCs to iECs and iVSMC. Each clone of hiPSCs was differentiated in multiple rounds, and low-SBP and high-SBP clones were always differentiated and analyzed in parallel. We performed RNA-seq analysis (n=6) and found that *SHOX2* was marginally significantly downregulated in iECs with the high-SBP allele of rs9833313 compared with the low-SBP allele (**Fig 7D**). Other neighboring genes *VEPH1, PTX3, SLC66A1L, RSRC1,* and *MLF1* were not differentially expressed. In iVSMCs, these local genes were not differentially expressed (**Fig 7D**). qPCR analysis in cells from more rounds of differentiation identified a significant, approximately 50% downregulation of *SHOX2* in iECs with the high-SBP allele of rs9833313 (n=24-25) and a significant but minor downregulation in iVSMCs (n=45) (**Fig 7E**). ChIP-qPCR analysis indicated a significant decrease of the enhancer histone marks H3K4me1 and H3K27Ac at the rs9833313 genomic region in iECs with the high-SBP allele of rs9833313 and no change in the binding of the inhibitory mark H3K27me3 (**Fig 7F**), consistent with a decrease of enhancer activity.

To identify genes and pathways linking SHOX2 to blood pressure regulation, we performed a GSEA analysis using the RNA-seq data mentioned above. At FDR < 10%, 22 gene sets were enriched in iECs with the low-SBP allele compared to the high-SBP allele, including several gene sets with potential relevance to blood pressure regulation (**Table S6**). No gene sets were enriched in iECs with the high-SBP allele or in iVSMCs with either allele. The 22 gene sets enriched in iECs with the low-SBP allele included 908 core enrichment genes. *SHOX2* was one of the core enrichment genes in a set of genes downregulated by KRAS activation. Two other core enrichment genes, GJA5 (Gap Junction Protein Alpha 5) in the myogenesis gene set and NKX2-5 (NK2 Homeobox 5) in a gene set upregulated by UV response, were among the 20 genes associated with SHOX2 directly or with one intermediate step based on a STRING analysis ^20^ (**Fig S14**). In addition, 30 of the 908 core enrichment genes were among the 250 blood pressure physiology genes that we previously curated ^7^. STRING analysis without restrictions on the number of intermediate steps showed that SHOX2 was associated with several of these blood pressure physiology genes via GJA5 (**Fig 7G**).

## Discussion

In this study, we applied a comprehensive multi-omics approach to unravel the epigenetic and genetic regulatory mechanisms governing gene expression in iECs and iVSMCs. By integrating DNA methylation, chromatin accessibility, chromatin interactions, and transcriptomic data, we characterized the unique epigenomic landscapes of these vascular cells and advanced our understanding of the regulatory architecture underlying vascular gene expression. Our findings underscore the importance of integrating multiple epigenetic modalities to understand the complex mechanisms of vascular gene regulations and identify potential targets for treating vascular disorders.

Compared to other epigenetic mechanisms, chromatin conformation has been substantially less studied in vascular cells ^21,22^. Notably, we have found that chromatin conformation plays a prominent role in the regulation of gene expression in vascular cells. This prominent role was suggested by a significantly strong correlation between chromatin interaction counts and gene expression levels and experimental results of the CTCF and RAD21 depletion study. CTCF and the cohesin complex function in complementary but distinct fashion ^23^. While CTCF and cohesin can cooperate to mediate the formation of chromatin loops, including topologically associated domains (TADs), not all CTCF-mediated loops depend on cohesin. In contrast, cohesin primarily stabilizes loops between enhancers and promoters, essential for sustained gene expression.

One of the new discoveries in our study is the identification of distinct epigenetic landscape in chromatin contact regions associated with specific regulatory interactions, such as PP, EP, and PT interactions. These regions displayed lower DNA methylation levels and greater chromatin accessibility than non-contact regions in both cell types, likely promoting transcription by facilitating transcription factor binding. In contrast, EE, TT, and ET interactions were characterized by higher DNA methylation, which may contribute to a more restrictive chromatin state. These findings reveal how different regulatory element interactions may influence gene expression in vascular cells through the coordination with other epigenetic features.

Most SNPs associated with complex traits are in noncoding genomic regions^7,8,24^. Understanding how these SNPs regulate gene expression and, thereby, influence their associated traits is one of the most important but difficult challenges in the genetic studies of complex traits ^25–27^. Our findings indicate that SNPs associated with vascular traits, such as blood pressure, stroke, coronary artery disease, and peripheral artery disease, are significantly more enriched in epigenetic features in iECs and iVSMCs, including chromatin loops and accessible regions, than SNPs associated with non-vascular traits. Furthermore, these SNPs are enriched in chromatin contacts involving PP, EP, and PT interactions. These exciting findings strongly suggest that SNPs influence gene expression through epigenetic mechanisms in a trait- and cell-type-dependent manner.

Our CRISPR-mediated editing of SNP rs9833313 provides a proof of principle demonstration of a vascular trait-associated SNP influencing the expression of a gene located hundreds of kbp away but within the same chromatin loops in vascular cells. The allelic sequences of rs9833313 affected the enhancer activity of the rs9833313 genomic region. The study highlights *SHOX2* as a gene of interest in BP regulation. *SHOX2* encodes a transcriptional factor important for the development of several organ systems including the skeleton and the sinoatrial and pacemaking development in the heart ^28,29^. Future studies should further investigate how SHOX2 specifically influences biological pathways underlying BP regulation and the therapeutic potential of these mechanisms.

In summary, this study provides an extensive map of the epigenetic landscape in iPSC-derived vascular cells, which highlights the importance of integrating multiple epigenetic mechanisms for understanding gene regulation. Targeted experiments revealed specific roles for nuclear architecture proteins and a BP-associated noncoding SNP in gene regulation in vascular cells, demonstrating the power for the epigenetic landscape to drive novel studies. We envision that the findings of this study will motivate numerous new lines of research on genetic and epigenetic regulation of vascular function and disease. Moreover, our approach of coupling integrated multi-omics analysis in trait-relevant cell types with targeted experimental interventions including precise genome editing in hiPSCs offers a valuable framework for investigating gene regulatory mechanisms in other cell types and disease contexts.

## Material and Methods

### hiPSCs and their differentiation into iECs and iVSMCs

The human iPSC line 039B was reprogrammed from urine cells and differentiated into Endothelial cells (iECs) and vascular smooth muscle cells (iVSMCs) as we described previously ^8,30,31^. iPSCs were cultured on 6-cm dishes coated with hESC-qualified Matrigel (Corning) using mTeSR™ Plus medium (STEMCELL Technologies). For differentiation, iPSCs were dissociated with Accutase (STEMCELL Technologies) and seeded onto Matrigel-coated 6-well plates at a density of 45,000–50,000 cells/cm² in mTeSR™ Plus medium supplemented with 10 µM Rock inhibitor Y-27632 (STEMCELL Technologies). After 24 hours, cells were treated with N2B27 medium (a 1:1 mixture of DMEM:F12 with Glutamax and Neurobasal media supplemented with N2 supplement and B27 supplement minus vitamin A; all from Life Technologies) containing 8 µM CHIR99021 (Selleck Chemicals) and 25 ng/ml BMP4 (PeproTech) for 3 days to generate mesoderm cells.

To induce endothelial lineage differentiation, mesoderm cells were cultured in StemPro-34 SFM medium (STEMCELL Technologies) supplemented with 200 ng/ml VEGF (PeproTech) and 2 µM forskolin (Abcam) for 2 days. On day 6, endothelial cells were purified using CD144 (VE-Cadherin) magnetic beads (Miltenyi Biotec). The purified CD144-positive cells were cultured in StemPro-34 SFM medium with 50 ng/ml VEGF for an additional 6 days before harvest.

For VSMC induction, mesoderm cells (on day 4) were cultured in N2B27 medium supplemented with 10 ng/ml PDGF-BB (PeproTech) and 2 ng/ml Activin A (PeproTech) for 2 days. Contractile VSMCs were subsequently induced by culturing in N2B27 medium containing 2 ng/ml Activin A and 2 µg/ml Heparin (STEMCELL Technologies) for 6 days. On day 12, VSMCs were enriched by depleting CD144-positive cells using CD144 magnetic beads. The purified VSMCs were replated and cultured in N2B27 medium supplemented with 2 ng/ml Activin A, 2 µg/ml Heparin, and 1 µM PD0325901 (a MEK inhibitor; Selleck Chemicals) for an additional 6 days before harvest.

### Genome editing in hiPSCs

The generation of isogenic hiPSCs with homozygous low-SBP and high-SBP alleles for the SNP rs9833313 was accomplished using a two-step genome editing approach, as we previously described ^8^. This process involved two key steps: deletion of the SNP rs9833313 region and homology-directed knock-in of the desired SNP alleles.

In the first step, two synthetic sgRNAs targeting a 50–200 bp region at the 5’ and 3’ ends of SNP rs9833313 were designed using the Cas-Designer web tool (http://www.rgenome.net/cas-designer/) for spCas9 and synthesized by Synthego (**Table S7**). The sgRNAs (3 µM each) were co-delivered with spCas9-2NLS Nuclease protein (3 µM) into hiPSCs using the Lipofectamine Stem Transfection Reagent (Invitrogen). After 48 hours, cells were dissociated with Accutase and subjected to single-cell sorting using the microfluidic-based Namocell HANA benchtop single-cell sorter (Bio-Techne) into 96-well plates. Single-cell clones were expanded for 10–14 days, and genomic DNA was extracted for initial characterization of the SNP deletion using PCR across the rs9833313 locus. Clones showing a shorter PCR fragment (∼185 bp deletion) were further confirmed by Sanger sequencing to validate the deletion of the SNP region.

In the second step, to reconstitute the SNP region with either the low-SBP rs9833313-A or high-SBP rs9833313-T allele, a homology-directed repair (HDR) approach was employed. A gRNA targeting the junction of the deleted rs9833313 region was designed (**Table S7**) and synthesized by Synthego. An ssDNA donor template (∼475 bp) containing the desired SNP allele and 200–300 bp homology arms on either side of rs9833313 was prepared. The donor DNA template was generated by amplifying the rs9833313 region from the parental iPSC line (039B) using Q5 High-Fidelity DNA Polymerase (NEB). The amplified product was cloned into the pBR322 plasmid, and the low-SBP rs9833313-A allele was confirmed by Sanger sequencing. The high-SBP rs9833313-T allele was introduced by site-directed mutagenesis using the QuikChange II Site-Directed Mutagenesis Kit (Agilent) and confirmed by sequencing. Donor ssDNA (∼475 bp) was prepared from the plasmids using the Guide-it Long ssDNA Production System (Takara Bio). The HDR template (500 ng ssDNA for either allele), along with two sgRNAs (3 µM each) and spCas9-2NLS Nuclease protein (3 µM), was transfected into hiPSCs containing the SNP region deletion using Lipofectamine Stem Transfection Reagent (Invitrogen). After 48 hours, single-cell cloning was performed using the Namocell HANA sorter, and clones were expanded for 10–14 days. Reconstitution of the SNP region was confirmed by PCR and subsequent Sanger sequencing of the amplified product. Clones with homozygous low-SBP rs9833313-A or high-SBP rs9833313-T alleles were further characterized for pluripotency, differentiation potential, and genetic stability by molecular karyotyping.

### Chromatin immunoprecipitation and qPCR

Chromatin immunoprecipitation (ChIP) was performed using antibodies for H3K4me1 (Abcam, ab8895), H3K27Ac (Millipore Sigma, MABE647), and H3K27me3 (Millipore Sigma, 07-449) and the EZ-Magna ChIP® G Chromatin Immunoprecipitation Kit (Millipore Sigma) according to the manufacturer’s instructions. Briefly, two million differentiated cells (iECs and iVSMCs) were washed with DPBS, crosslinked with 1% formaldehyde for 10 minutes at room temperature, and quenched with 10X glycine for 5 minutes. The cells were washed twice with cold PBS containing 1X Protease Inhibitor Cocktail II (Millipore Sigma) and lysed with cell lysis buffer containing 1X Protease Inhibitor Cocktail II for 15 minutes on ice, with brief vortexing every 5 minutes.

The nuclear fraction was collected as a pellet by centrifugation at 800 × g for 5 minutes at 4°C and resuspended in nucleus lysis buffer containing 1X Protease Inhibitor Cocktail II. The lysate was sonicated using a Covaris S220 focused-ultrasonicator at 4°C to generate DNA fragments of 200–800 bp. The sonicated solution was centrifuged at 12,000 × g to remove insoluble material. The supernatant containing sheared, crosslinked chromatin was diluted 10-fold in dilution buffer containing Protease Inhibitor Cocktail II, and ∼5% of the supernatant was saved at 4°C as the “Input” control. The remaining supernatant was incubated overnight with anti-H3K4me1 (2µg), anti-H3K27Ac (2µg), anti-H3K27me3 (2µg), or negative control anti-rabbit IgG (2mg), and 20 µl of fully suspended Protein G magnetic beads.

Using a magnetic separator, the Protein G magnetic bead–protein–DNA complexes were separated and washed sequentially with cold buffers, including low salt immune complex wash buffer, high salt immune complex wash buffer, LiCl immune complex wash buffer, and TE buffer, for 5 minutes each. The protein/DNA complexes were eluted with ChIP elution buffer supplemented with Proteinase K. The elution mixture was incubated at 62°C for 2 hours with shaking, followed by 95°C for 10 minutes to reverse crosslinking.

The Protein G magnetic beads were separated using a magnet, and the supernatant was collected for DNA purification using a spin filter. The purified DNA was eluted from the spin filter using elution buffer and analyzed by qPCR using primers specific to the SNP rs9833313 region (**Table S7**).

### Lentiviral-mediated knockdown of CTCF and RAD21

293T cells were transiently transfected with the packaging vectors VSV-G and pΔ8.9 along with lentiviral vectors to knock down CTCF, RAD21, or the control vector, pLKO.1 puro (#8453 from Addgene). Two different shRNA lentiviral constructs were used for each target gene. To knockdown CTCF, we used TRCN0000218498, and TRCN0000014548 shRNA constructs from Millipore Sigma. For the RAD21 knockdown, we used TRCN0000148110, and TRCN0000281202 shRNA constructs from Millipore Sigma (**Table S7**). Viral media was collected 48 and 72 hrs post-infection and spun at 6000g overnight at 4°C. The concentrated bottom 500 μl of the viral solution was used for infection. iECs were seeded in a Matrigel-coated six-well plate, and 50 μl of the concentrated viral solution was added to each well. The next day, cells were fed with new media. After 48 hrs of infection, cells infected with the virus were selected using 1 μg/ml of Puromycin for the next 48 hrs.

### Quantitative real-time PCR

Total RNA was extracted from cell samples using TRIzol reagent (Life Technologies) and converted to cDNA using random hexamer primers and the RevertAid First Strand cDNA Synthesis Kit (Thermo Scientific). Quantitative real-time PCR (qPCR) analysis for mRNA expression was performed as previously described ^32^. Primer sequences are listed in **Table S7**. mRNA expression levels were normalized to the endogenous control GAPDH or 18S rRNA using the comparative Ct (ΔΔCt) method.

### Western blot analysis

Whole-cell lysates were prepared by resuspending the 1×10^6^ viral-infected iECs in 100 mL Laemmli buffer and boiling for 10 min. An equal amount of lysates per sample was loaded into a pre-cast 8-16% gradient Mini PROTEAN TGX gels (Biorad, 4561105), and proteins were transferred to the nitrocellulose membrane. After blocking in 5% milk in 1X PBS for 30 mins, blots were probed with the anti-b-actin (Abcam, ab219733, 1:10,000), anti-CTCF (Millipore-Sigma 07-729), and anti-RAD21 (Abcam, ab992, 1:1000) for overnight at 4°C. Blots were washed with the tris-buffered saline with Tween-20 (TBST, Boston Bioproducts, IBB 180) thrice (10 mins each wash) and probed with the secondary antibody (1:6000 anti-rabbit HRP, Abcam, ab205718) for one hour at room temperature. After washing thrice with TBST, blots were developed using Immobilon Western Chemiluminescent HRP Substrate (Millipore Sigma, WBKLS0500) and imaged on an Amersham Imager 680. Information on antibody was listed in **Table S8**.

### RNA sequencing

Total RNA was extracted from cell samples using the TRIzol reagent (Invitrogen, Carlsbad, CA, USA). RNA quality assessments were conducted, including 1% Agarose gel electrophoresis for visualizing RNA integrity, Nanodrop readings to quantify RNA amount and assess purity, and Agilent 2100 Bioanalyzer (Agilent Technologies, Palo Alto, CA, USA) for further RNA integrity evaluation. Before cDNA library construction, approximately 1 μg of total RNA underwent rRNA depletion using the NEBNext rRNA Depletion Kit (NEB, Ipswich, MA, USA, Cat# E6318). RNA-seq libraries were then prepared following the whole transcriptome sequencing protocol of Novogene (https://www.novogene.com/us-en/resources/blog/a-basic-guide-to-rna-sequencing/). These multiplexed libraries were subsequently sequenced on a NovaSeq 6000 sequencer (Illumina, San Diego, CA, USA) with paired-end reads of 2 × 150 bp, generating approximately 86 million reads for each library (**Table S1** and **S5**).

### RNA-seq data analysis

RNA-seq data was analyzed as we described ^33,34^. Raw RNA-Seq sequence reads were pre-processed using Trim Galore (v0.6.2) (http://www.bioinformatics.babraham.ac.uk/projects/trim_galore/). Adapters and reads with low quality (base quality < 20) were removed prior to further analysis. Reads longer than 20bp after the trimming were aligned to the human genome (hg38) using hisat2 (v2.2.1) ^35^. Transcript assembly and quantification were performed using StringTie2 (v2.2.1) ^36^, based on the GENCODE reference annotation (v38).

### Differential expression analysis

Differentially expressed genes (DEGs) between two groups of iPSC-derived cells were detected using DESeq2 ^37^. For knockdown experiments, pLKO-infected iECs served as controls for shCTCF- and shRad21-infected iECs, respectively. Gene count data obtained from StringTie2 were used as input for DESeq2 analysis. A false discovery rate (FDR) < 0.05 was applied to define significant differential expression.

### Reduced representation bisulfite sequencing (RRBS)

Genomic DNA of iECs and iVSMCs was extracted and processed following Diagenode’s Premium RRBS protocol (https://www.diagenode.com/). In brief, genomic DNA was digested using Msp1, followed by end-repair. Next, premium Methyl Unique Dual Indexes (UDI) and Unique Molecular Identifiers (UMI) Adapters were ligated to the resulting fragments. Then, UDI-UMI-tagged fragments are size-selected to enrich CpG-rich regions with gene regulatory potential. Finally, the enriched DNA underwent bisulfite conversion, followed by sequencing on an Illumina Novaseq 6000 platform with paired-end 50 bp reads (**Table S2**).

### RRBS data analysis

RRBS data was analyzed as we described ^38^. Bisulfite-converted reads were processed using Trim Galore with non-directional and paired setting to remove adapters and low-quality sequences. The trimmed reads were aligned to the human genome (hg38) using bismark (v0.23.1dev) ^39^. Next, PCR duplicates were removed by identifying and collapsing reads with identical UMI tags. Then, Bismark_methylation_extractor was employed to extract methylation calls for individual CpG sites. Bisulfite conversion efficiency was assessed using methylated and unmethylated spike-in controls. Finally, for each CpG site, DNA methylation rate was calculated as the percentage of unconverted cytosines within each RRBS library. To visualize global methylation profiles in these iPSC-derived cells, the top 1% of most variable CpG sites across all samples were selected for unsupervised hierarchical clustering and principal component analysis (PCA). Promoter is defined as 2kbp upstream and 200bp downstream of the transcription start site (TSS).

### Detection of differentially methylated regions

DMRs were identified using Metilene, a preferred tool specifically designed for RRBS data ^40^. Metilene applies a binary segmentation algorithm combined with a two-dimensional Kolmogorov–Smirnov test to de novo identify genomic regions that maximize methylation differences between two groups. For this analysis, a minimum mean methylation difference between groups of 0.05 and a minimum of five CpGs within a region were required. Statistical significance of DMRs was determined using an FDR threshold of < 0.05.

### ATAC sequencing

Genomic DNA was extracted from these iPSC-derived cells to prepare ATAC-seq libraries. The OMNI ATAC-seq protocol ^41^ was adopted for isolating nuclei and applying transposase reaction. Following the reaction, the libraries were purified using two-sided SPRI beads selection to isolate DNA fragments within the 100-500bp range. The final multiplexed ATAC-seq libraries were sequenced on an Illumina Novaseq 6000 platform with paired-end 150 bp reads (**Table S3**).

### ATAC-seq analysis

The analysis of ATAC-seq data adhered to the ENCODE Standards and Processing Pipeline for ATAC-seq, accessible at https://www.encodeproject.org/atac-seq/, which is designed for automated end-to-end quality control and processing of ATAC-seq. In brief, raw ATAC-seq reads were quality-trimmed and aligned to the human genome (hg38) using Burrows-Wheeler Aligner ^42^. Peak calling was performed using MACS2 from the alignment files to identify regions of open chromatin ^43^. Processed output files, including bigWig format for genome browser, were generated. Statistically significant ATAC-seq peaks, indicative of open chromatin, were discerned based on a q-values threshold < 0.05.

### Micro-C library preparation

Micro-C assay ^44^ was performed using Dovetail® Micro-C Kit following the manufacturer’s User Guide and as we described ^8,45^. Briefly, up to 1M cells were harvested and pellet were frozen at -80°C for at least 30 minutes. iECs and iVSMCs were sequentially crosslinked with disuccinimidyl glutarate and formaldehyde to preserve chromatin interactions. Genomic DNA was digested in situ with MNase Enzyme mix to achieve 40%-70% mononucleosomes digestion. The DNA was then released from the cells and put through a sequential process of binding to chromatin capture beads, end polishing, bridge ligation, intra-aggregate ligation, crosslink reversal, and DNA purification. Library was made from the purified DNA and sequenced on Novaseq 6000 platform, generating approximately 800 million 150bp paired-end reads (**Table S4**).

### Micro-C data analysis

Micro-C data was analyzed as we described ^8,45^. Paired-end sequencing reads were processed with Trim Galore to remove adaptor and low-quality reads. Processed reads were analyzed using JuicerBox, a streamline pipeline for processing HiC or Micro-C data, to generate hic file for direct visualization and topolically associating domain (TAD) calling using Arrowhead ^46^. Additionally, Dovetail Micro-C Data Processing Guide was followed to generate .mcool file, which were used for loop calling with Mustache ^47^.

### Correlation between gene expression and epigenetic features

The Spearman correlation between gene expression levels (log2(FPKM+1)) and chromatin accessibility at gene promoters was assessed. For promoters with multiple ATAC-seq peaks, the peak with the highest signal was used for correlation analysis. To evaluate the relationship between gene expression and DNA methylation in promoter/enhancer regions, we defined promoters as the region spanning -200 bp to +2 kb from the transcription start site (TSS). Enhancer regions were identified based on genomic positions obtained from the Ensembl Regulatory Bulid (hg38). Additionally, number of chromatin interactions within genes were counted from micro-C data and their correlation with gene expression were analyzed.

### Regulatory elements in chromatin contact regions

Chromatin contact regions form DNA loops, allowing distant regulatory elements to interact. Regulatory elements within these chromatin contacts regions, including promoters, enhancers, and transcription factor binding sites (TFBS), were annotated using the Ensembl Regulatory Build (hg38). By focusing on the elements within the chromatin contacts, we categorized the regulatory interactions into six distinct classes based on the type of regulatory elements involved: promoter-promoter (PP), enhancer-enhancer (EE), TFBS-TFBS (TT), enhancer-promoter (EP), enhancer-TFBS (ET), and promoter-TFBS (PT) interactions.

### Characterization of interactions between regulatory elements

Several key epigenomic features were characterized within chromatin contact regions. Specifically, we examined the DNA methylation levels, chromatin accessibility (as measured by signal value of ATAC-seq peak), the presence of target SNPs, and the gene expression levels associated with these regulatory interactions. The expression of genes within 5 kbp upstream and downstream of the regulatory element were compared among different regulatory interaction types. Difference in these epigenetic features and gene expression were determined using the two-sample Wilcoxon test.

### Impact of Rad21 or CTCF knockdown on genes within chromatin loops

To explore whether Rad21 or CTCF regulates downstream targets through chromatin interactions, we categorized genes into two groups: those located within chromatin loops and those outside them. DEGs were then counted in each group following Rad21 or CTCF knockdown, generating a 2×2 contingency table. A Fisher’s exact test was performed on this contingency table, and odds ratios were calculated to assess the association between chromatin loops and DEGs upon knockdown of Rad21 or CTCF. Additionally, the relationship between chromatin contacts, Rad21/CTCF genome occupancies, and DEGs was assessed using the same approach.

### GSEA Analysis

Gene Set Enrichment Analysis (GSEA, version 4.3.3) ^48^was performed using normalized count data for all genes from CTCF- and RAD21-depleted iECs versus empty vector-infected iEC controls. The analysis utilized the Hallmark subset of canonical pathways (h.all.v2024.1.Hs) with 1000 permutations, and the permutation type was set to “gene_set.” Gene sets with a false discovery rate (FDR) < 0.05 were considered significant.

### Enrichment analysis of gene ontology and pathways

Gene ontology (GO) term and pathway enrichment of the genes of interests were performed using clusterProfiler ^49^.

### Statistical analysis

Quantitative variables, such as gene expression and epigenetic features, are presented as mean ± standard error (SEM). Categorical variables, including SNPs and regulatory interactions, are reported as proportions. Differences in quantitative variables between groups were assessed using the Wilcoxon signed-rank test, while differences in categorical variables were evaluated using a two-sample proportion test. SNP enrichment within specific omics datasets was analyzed using hypergeometric tests. The Kolmogorov-Smirnov test was used to evaluate differences in the probability density of gene expression across groups. All statistical analyses were performed using the R statistical software package (v4.2.2).

## Author contributions

PL analyzed data. AR performed the knockdown experiment. YL led the omics assays and analyzed data. MKM performed the rs9833313 experiment. QQ analyzed data. RP, BT, and CS performed experiments. JH and JR contributed to data analysis. XB, ASG, AWC, and AMG contributed to study design or data interpretation. PL, SR, and ML led the study. PL, QQ, SR, and ML drafted the manuscript. AR, YL, MKM, and RP contributed to the drafting of the manuscript. All authors edited and approved the manuscript.

