## Supplementary figures for "Global and genetic regulation of gene expression in human endothelial and vascular smooth muscle cells"

[1]Department of Physiology, University of Arizona College of Medicine – Tucson, Tucson, AZ, USA; [2]Molecular Systems Medicine Initiative, University of Arizona Health Sciences, Tucson, AZ, USA; [3]Versiti Blood Research Institute, Milwaukee, WI, USA; [4]Department of Respiratory Medicine, Sir Run Run Shaw Hospital and Institute of Translational Medicine, Zhejiang University School of Medicine, Hangzhou, Zhejiang China; [5]Department of Cell Biology, Neurobiology & Anatomy, Medical College of Wisconsin, Milwaukee, WI, USA; [6]The Jackson Laboratory, Bar Harbor, ME, USA; [7]Department of Physiology, Medical College of Wisconsin, Milwaukee, WI, USA; [8]Department of Pediatrics, Division of Hematology, Oncology, and Transplantation, Medical College of Wisconsin, Milwaukee, WI, USA

*co-first authors.

[#]co-corresponding authors:

Pengyuan Liu,

Sridhar Rao,

Mingyu Liang,

### Supplementary figure legends

**Figure S1. RNA-seq analysis of iEC and iVSMC. Related to Figure 1.**
(**A**) Heatmap of RNA-seq data using all gene expressions.
(**B**) PCA (Principal Component Analysis) of RNA-seq data using all gene expressions.
(**C, D**) Gene Ontology (GO) terms of genes enriched in iEC (C) and iVSMC (D).
(**E, F**) Pathways of genes enriched in iEC (E) and iVSMC (F).

**Figure S2. DNA methylation analysis of iECs and iVSMCs. Related to Figure 1.**
(**A, B**) Heatmap (A) and PCA (B) of RRBS data based on the top 10,000 CpG sites that varied most across iECs and iVSMCs.
(**C**) Frequency of DMRs with different methylation differences between iECs and iVSMCs. DMRs, differentially methylated regions.
(**D**) Volcano plot showing DMRs between iEC and iVSMC. Methylation difference represents the mean difference in promoter methylation at DMRs between iECs and iVSMCs.
(**E**) Genomic annotation of DMRs identified between iECs and iVSMCs.

**Figure S3. Distinct chromatin opening patterns between iECs and iVSMCs. Related to Figure 1.**
(**A**) Peak profiles of iEC and iVSMC. TSS, transcript start site.
(**B**) Average profile of ATAC-seq chromatin opening binding to TSS region in iECs and iVSMCs.
(**C**) Genomic annotation of chromatin opening in iECs and iVSMCs.

**Figure S4. Number of DNA loops and distribution of distance between chromatin contacts in iECs and iVSMCs. Related to Figure 1.**
(**A**) Venn diagrams showing number of DNA loops detected at 4kb and 16kb resolution.
(**B**) Distribution of distance between chromatin contacts.
DNA loops detected at 4kb, 8kb and 16kb resolution were analyzed separately.

**Figure S5. Genomic annotation of chromatin contact regions in iECs and iVSMCs. Related to Figure 1.**
Chromatin contacts of DNA loops were detected at 4kb, 8kb and 16kb resolution, respectively.

**Figure S6. Visualization of epigenomic landscapes of iEC and iVSMC for selected marker genes of EC and VSMC. Related to Figure 2.**
Genomic track show interaction loops, TADs, chromatin accessibility (ATAC_seq), DNA methylation (RRBS), gene expression (RNA_seq), and super-enhancers from dbSUPER for AGTR1 (A), NOS3 (B), and ACE (C).
TAD, topologically associating domain; HUVEC: Human umbilical vein endothelial cell; SE, super-enhancer.
Y-axis scale between cell type are identical for direct comparison.

**Figure S7. Genes with high expression levels tend to exhibit greater chromatin accessibility. Related to Figure 3.**
**(A, B)** Heatmap and average profile of ATAC-seq chromatin accessibility at the transcription start site (TSS) regions for different gene expression groups in iECs (**A**) and iVSMCs (**B**).
Genes were classified into four groups based on expression quantiles: Q1 (<25%), Q2 (25–50%), Q3 (50–75%), and Q4 (>75%).

**Figure S10. Number of different types of regulatory elements within chromatin contacts. Related to Figure 4.**
E, enhancer; P, promoter; T, transcription factor binding site. Chromatin contacts of DNA loops were detected at 4kb, 8kb and 16kb resolution, respectively.

**Figure S11. CTCF and RAD21 regulate gene expression in a chromatin organization-dependent manner. Related to Figure 5.**
(**A**) Knockdown efficiency of CTCF and RAD21 in ECs, as determined by qPCR. N=3 per shRNA; two-sided t tests were used to examine expression difference between groups; *, $p<0.05$.
(**B**) GO/pathway enrichment analysis of DEGs in iECs upon knockdown of Rad21 or CTCF.

**Figure S12. CTCF and RAD21 regulate gene expression in a chromatin organization-dependent manner. Related to Figure 5, but showing chromatin contacts at 4kb or 16kb resolution instead of the 8kb resolution shown in Figure 5.**
(**A, B**) Genes located in regulatory regions (such as DNA loops, chromatin contact, or near RAD21 or CTCF occupancy sites) are more likely to be altered in iECs, particularly following RAD21 knockdown. Overlapping represent overlapped regions between chromatin contacts and occupancy binding sites of CTCF or RAD21.
(**C, D**) Genes near chromatin contacts involving regulatory interactions are more likely to be altered in response to CTCF or RAD21 knockdown in iECs.

**Figure S13. Two-step genome editing scheme and confirmation of genomic deletion in Step 1. Related to Figure 7.**
(**A**) Pipeline for prioritizing BP-related SNPs for the the proof of principle study.
(**B**) Schematic diagram showing the two-step editing approach (deletion and re-constitution) for generating isogenic hiPSC lines with homozygous allele at SNP rs9833313.
(**C**) Location and size of ssDNA donor and across-region PCR amplicons at rs9833313.
(**D**) Across-region PCR confirmed the deletion of the SNP region, shown as shorter amplicon.
(**E**) rs9833313 region deletion confirmed by Sanger sequencing.

**Figure S14. Genes associated with SHOX2 based on a STRING analysis. Related to Figure 7.**
Genes associated with SHOX2 directly or at one step away were shown. Colors of nodes indicate functional clusters. Edges within a cluster are shown as solid lines, and edges between clusters are shown as dotted lines.

### Supplementary tables (submitted as an Excel file)

**Supplementary Table S1**. Quality control metrics for RNA-seq libraries.

**Supplementary Table S2**. Quality control metrics for RRBS libraries.

**Supplementary Table S3**. Quality control metrics for ATAC-seq libraries.

**Supplementary Table S4**. Quality control metrics for micro-C libraries.

**Supplementary Table S5**. QC metrics of RNA-seq libraries with knockdown of CTCF or Rad21.

**Supplementary Table S6**. Gene sets enriched in iECs with the low-SBP allele of rs9833313.

**Supplementary Table S7**. Sequences of primers, shRNA, and sgRNA used in the study.

**Supplementary Table S8**. List of antibodies used.

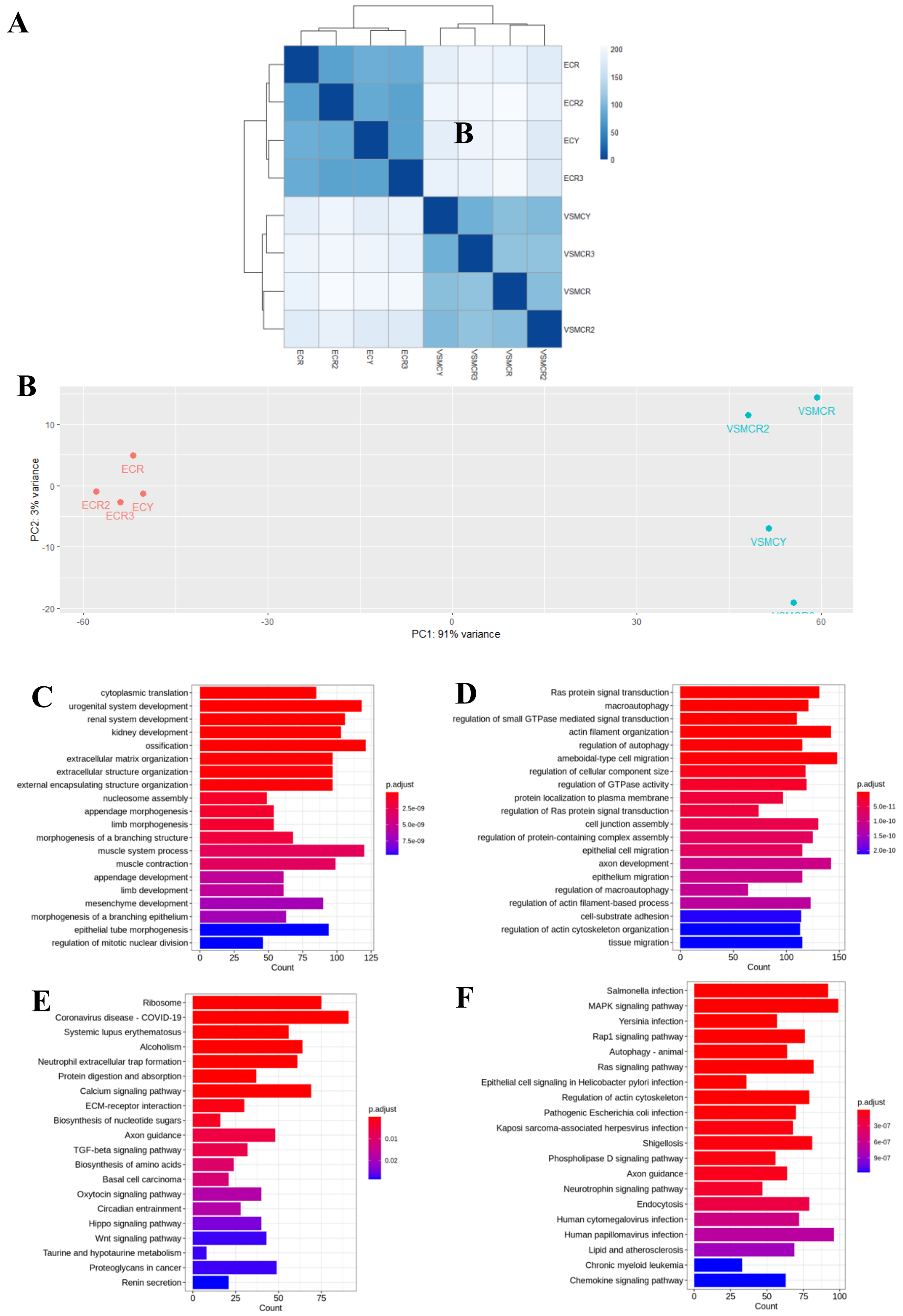


**Figure S1. RNA-seq analysis of iEC and iVSMC. Related to Figure 1.**
(**A**) Heatmap of RNA-seq data using all gene expressions.
(**B**) PCA (Principal Component Analysis) of RNA-seq data using all gene expressions.
(**C, D**) Gene Ontology (GO) terms of genes enriched in iEC (C) and iVSMC (D).
(**E, F**) Pathways of genes enriched in iEC (E) and iVSMC (F).

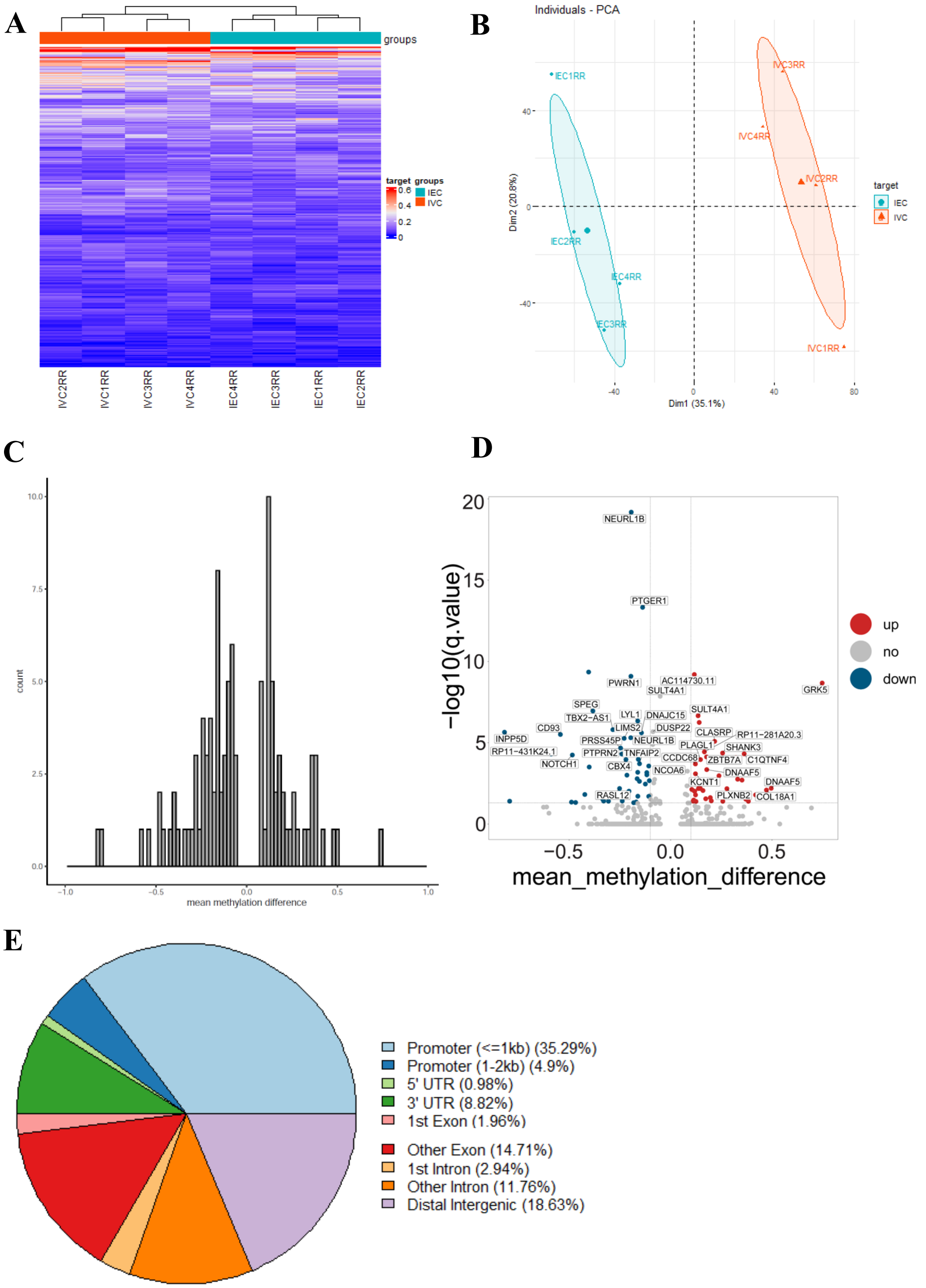


**Figure S2. DNA methylation analysis of iECs and iVSMCs. Related to Figure 1.**

(**A, B**) Heatmap (A) and PCA (B) of RRBS data based on the top 10,000 CpG sites that varied most across iECs and iVSMCs.

(**C**) Frequency of DMRs with different methylation differences between iECs and iVSMCs. DMRs, differentially methylated regions.

(**D**) Volcano plot showing DMRs between iEC and iVSMC. Methylation difference represents the mean difference in promoter methylation at DMRs between iECs and iVSMCs.

(**E**) Genomic annotation of DMRs identified between iECs and iVSMCs.

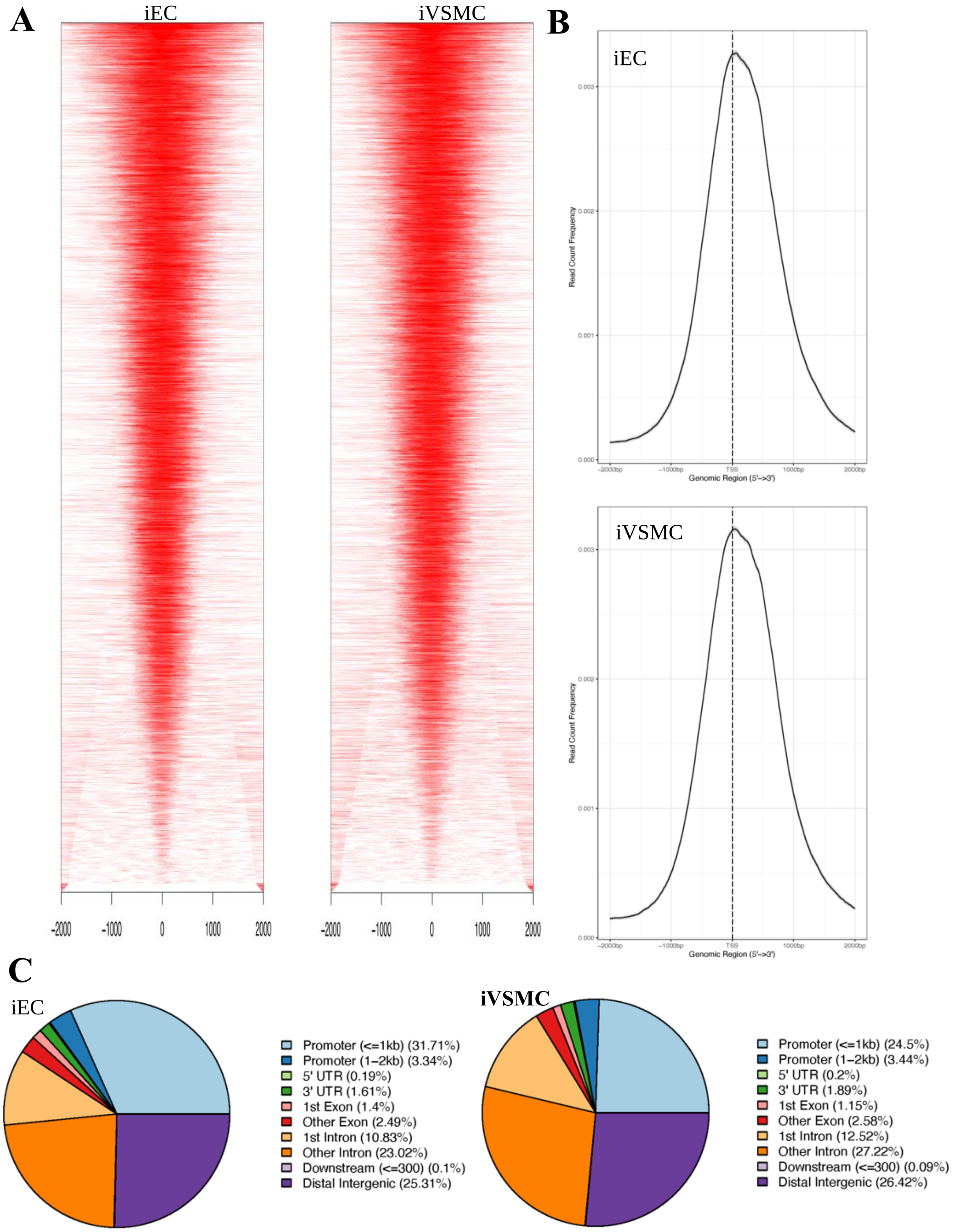


**Figure S3. Distinct chromatin opening patterns between iECs and iVSMCs. Related to Figure 1.**
(**A**) Peak profiles of iEC and iVSMC. TSS, transcript start site.
(**B**) Average profile of ATAC-seq chromatin opening binding to TSS region in iECs and iVSMCs.
(**C**) Genomic annotation of chromatin opening in iECs and iVSMCs.

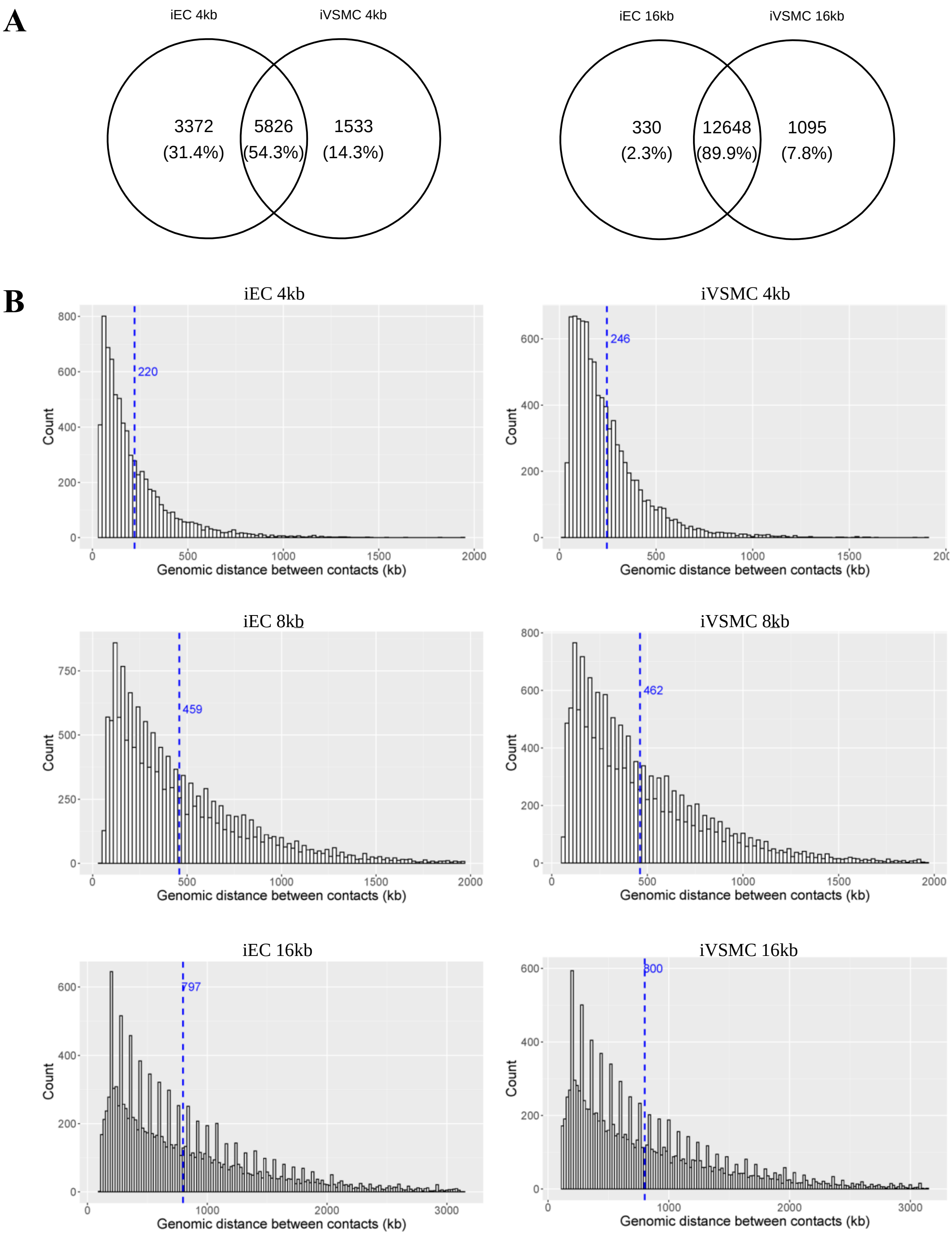


**Figure S4. Number of DNA loops and distribution of distance between chromatin contacts in iECs and iVSMCs. Related to Figure 1.**

(**A**) Venn diagrams showing number of DNA loops detected at 4kb and 16kb resolution.

(**B**) Distribution of distance between chromatin contacts.

DNA loops detected at 4kb, 8kb and 16kb resolution were analyzed separately.

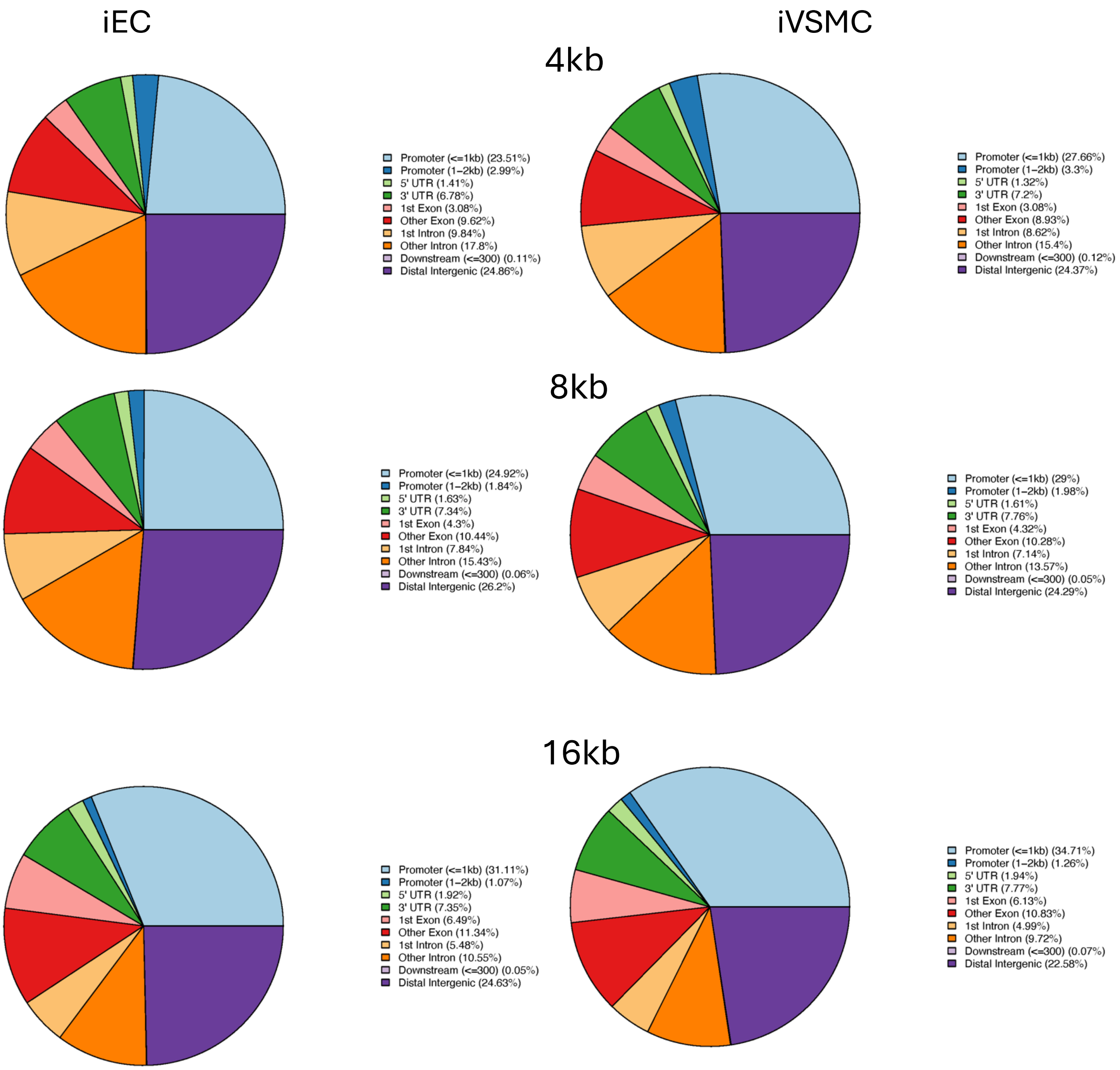


**Figure S5. Genomic annotation of chromatin contact regions in iECs and iVSMCs. Related to Figure 1.**

Chromatin contacts of DNA loops were detected at 4kb, 8kb and 16kb resolution, respectively.

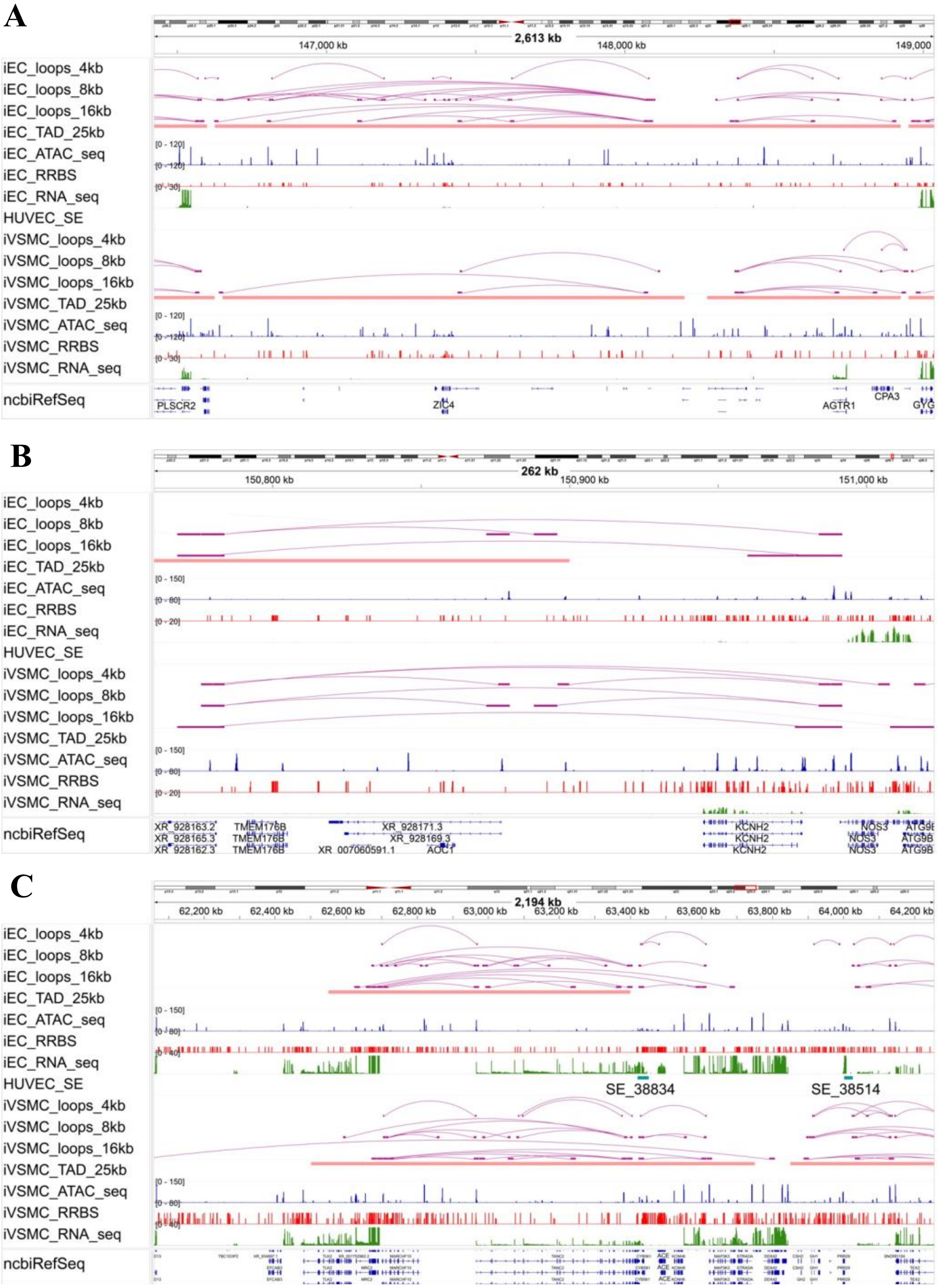


**Figure S6. Visualization of epigenomic landscapes of iEC and iVSMC for selected marker genes of EC and VSMC. Related to Figure 2.**

Genomic track show interaction loops, TADs, chromatin accessibility (ATAC_seq), DNA methylation (RRBS), gene expression (RNA_seq), and super-enhancers from dbSUPER for AGTR1 (A), NOS3 (B), and ACE (C).

TAD, topologically associating domain; HUVEC: Human umbilical vein endothelial cell; SE, super-enhancer.

Y-axis scale is identical between cell types for direct comparison.

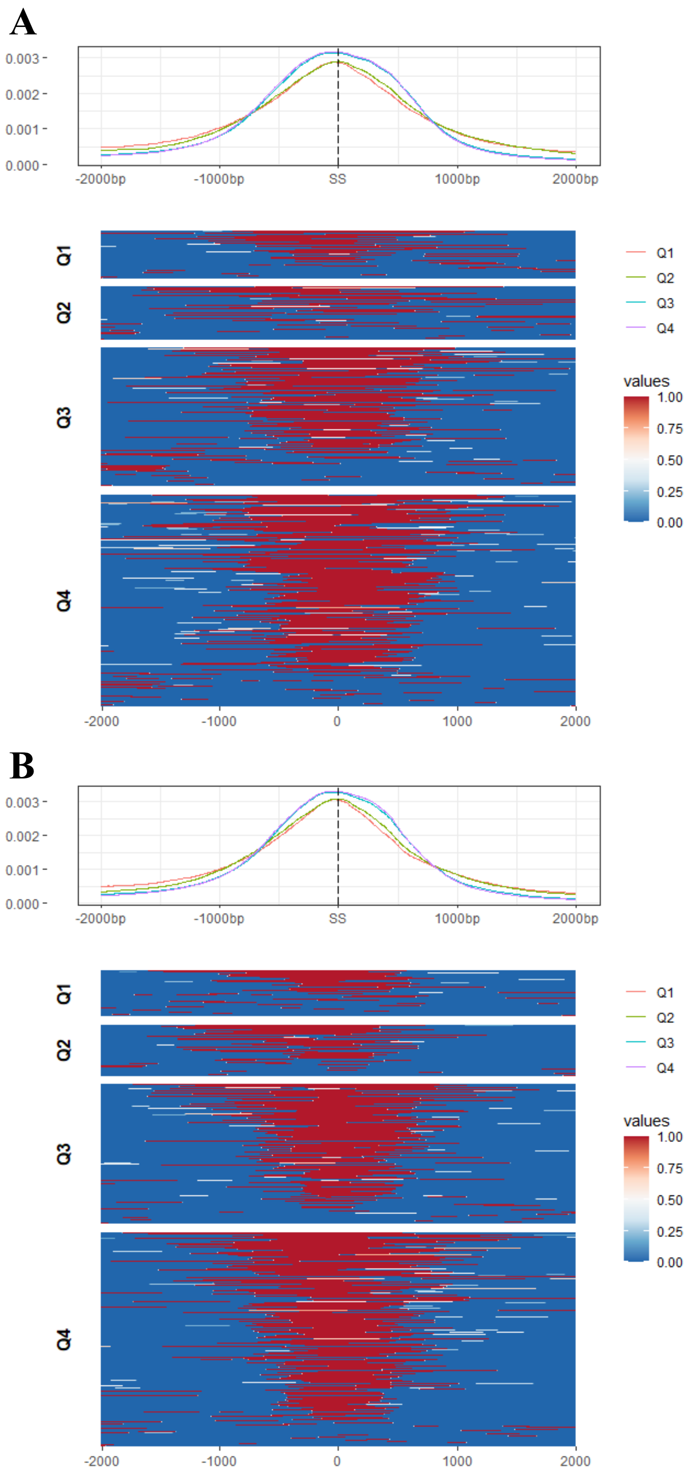


**Figure S7. Genes with high expression levels tend to exhibit greater chromatin accessibility. Related to Figure 3.**

**(A, B)** Heatmap and average profile of ATAC-seq chromatin accessibility at the transcription start site (TSS) regions for different gene expression groups in iECs (**A**) and iVSMCs (**B**).

Genes were classified into four groups based on expression quantiles: Q1 (<25%), Q2 (25–50%), Q3 (50–75%), and Q4 (>75%).

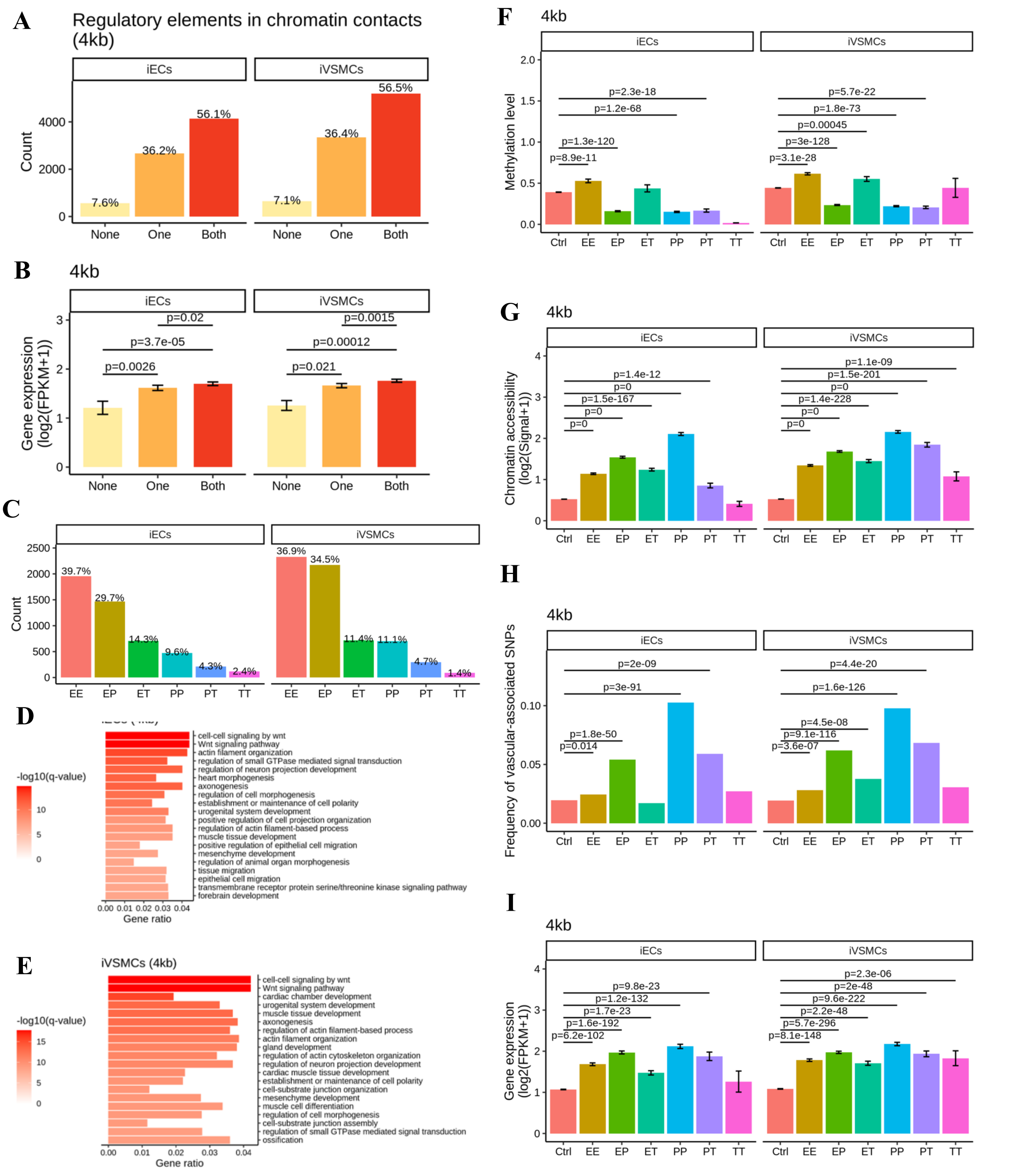


**Figure S8. Chromatin contact regions (4kb) in iECs and iVSMCs: regulatory element interactions, epigenetic features, and gene expression. Related to Figure 4.**

Chromatin contacts were detected at 4kb resolution.

(**A**) Chromatin contact regions of DNA loops were classified based on the presence of regulatory elements: "None" indicates that neither contact region of a loop contains a regulatory element; "One"

indicates that one of the contact regions of a loop contains a regulatory element; and "Both" indicates that both contact regions of a loop contain regulatory elements.
(**B**) Expression levels of genes located near chromatin contact regions with varying configurations of regulatory elements.
(**C**) Distribution of interaction types between regulatory elements within chromatin contact regions. Key interaction types include promoter-promoter (PP), enhancer-promoter (EP), enhancer-enhancer (EE), enhancer-transcription factor binding site (ET), promoter-transcription factor binding site (PT), and transcription factor binding site-transcription factor binding site (TT) interactions.
(**D, E**) GO and pathway enrichment analysis of genes near chromatin contact regions in iECs (D) and iVSMCs (E).
(**F**) DNA methylation levels in chromatin contact regions with different types of regulatory interactions.
(**G**) Chromatin accessibility in chromatin contact regions associated with various regulatory interactions.
(**H**) Enrichment of target SNPs in chromatin contact regions with different types of regulatory interactions.
(**I**) Expression levels of genes near chromatin contact regions with distinct regulatory element interactions.
Gene expression and epigenetic features are presented as mean ± standard error. Differences in quantitative variable between groups were assessed using Wilcoxon signed rank test, while differences in SNP proportions were evaluated using a two-sample proportion test.

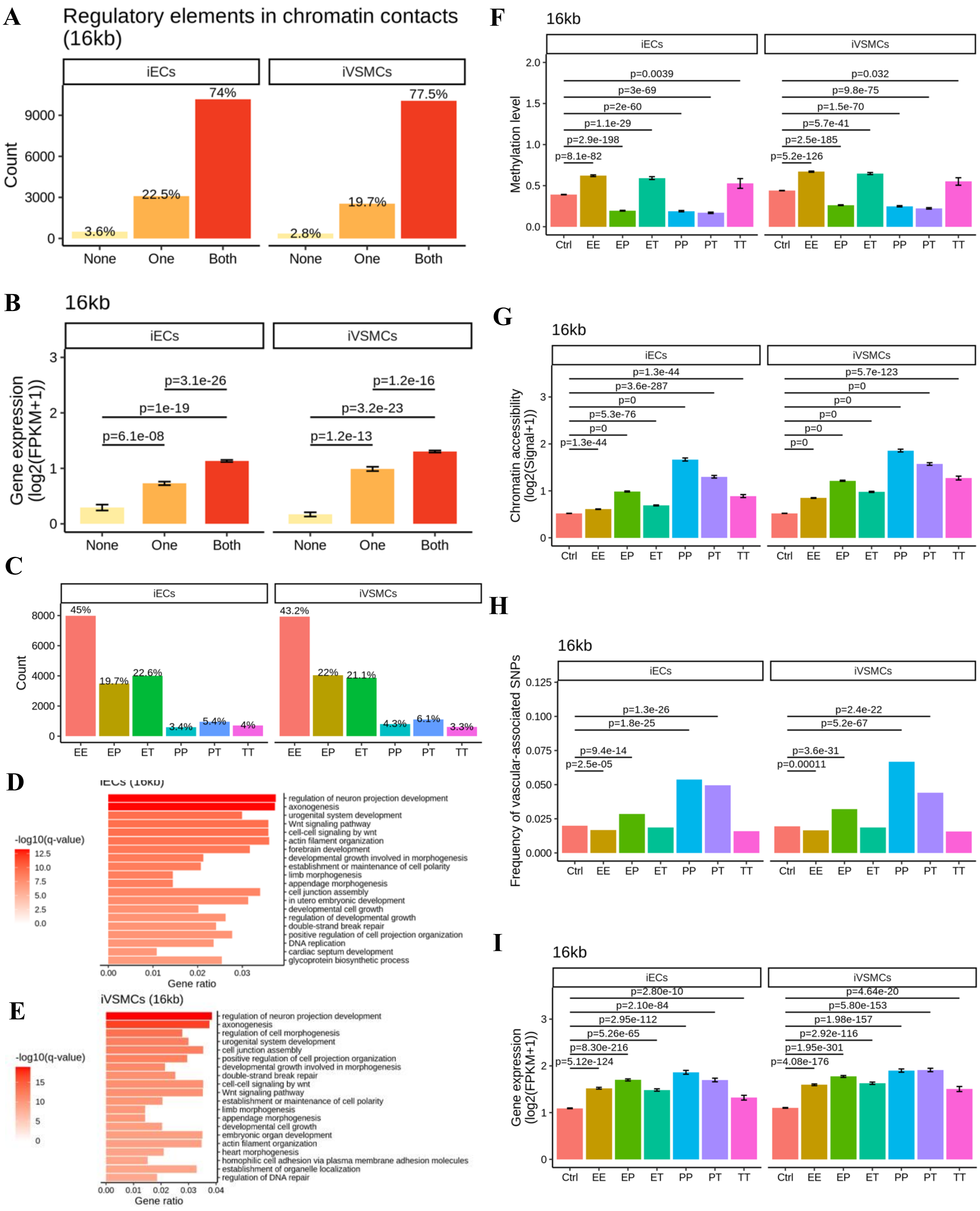


**Figure S9. Chromatin contact regions (16kb) in iECs and iVSMCs: regulatory element interactions, epigenetic features, and gene expression. Related to Figure 4.**
Chromatin contacts were detected at 16kb resolution.

(**A**) Chromatin contact regions of DNA loops were classified based on the presence of regulatory elements: "None" indicates that neither contact region of a loop contains a regulatory element; "One" indicates that one of the contact regions of a loop contains a regulatory element; and "Both" indicates that both contact regions of a loop contain regulatory elements.
(**B**) Expression levels of genes located near chromatin contact regions with varying configurations of regulatory elements.
(**C**) Distribution of interaction types between regulatory elements within chromatin contact regions. Key interaction types include promoter-promoter (PP), enhancer-promoter (EP), enhancer-enhancer (EE), enhancer-transcription factor binding site (ET), promoter-transcription factor binding site (PT), and transcription factor binding site-transcription factor binding site (TT) interactions.
(**D, E**) GO and pathway enrichment analysis of genes near chromatin contact regions in iECs (D) and iVSMCs (E).
(**F**) DNA methylation levels in chromatin contact regions with different types of regulatory interactions.
(**G**) Chromatin accessibility in chromatin contact regions associated with various regulatory interactions.
(**H**) Enrichment of target SNPs in chromatin contact regions with different types of regulatory interactions.
(**I**) Expression levels of genes near chromatin contact regions with distinct regulatory element interactions.
Gene expression and epigenetic features are presented as mean ± standard error. Differences in quantitative variable between groups were assessed using Wilcoxon signed rank test, while differences in SNP proportions were evaluated using a two-sample proportion test.

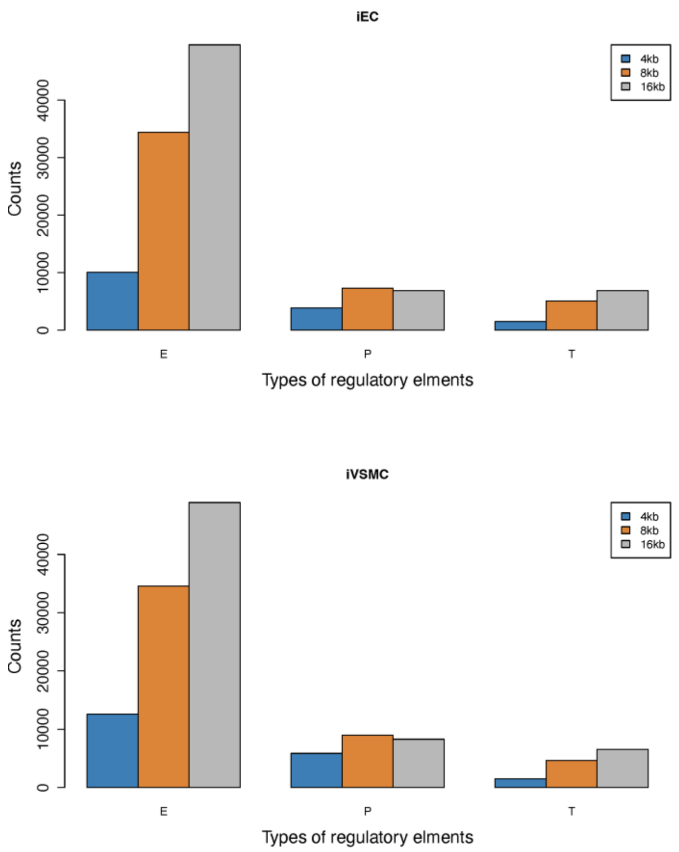


**Figure S10. Number of different types of regulatory elements within chromatin contacts. Related to Figure 4.**
E, enhancer; P, promoter; T, transcription factor binding site. Chromatin contacts of DNA loops were detected at 4kb, 8kb and 16kb resolution, respectively.

A

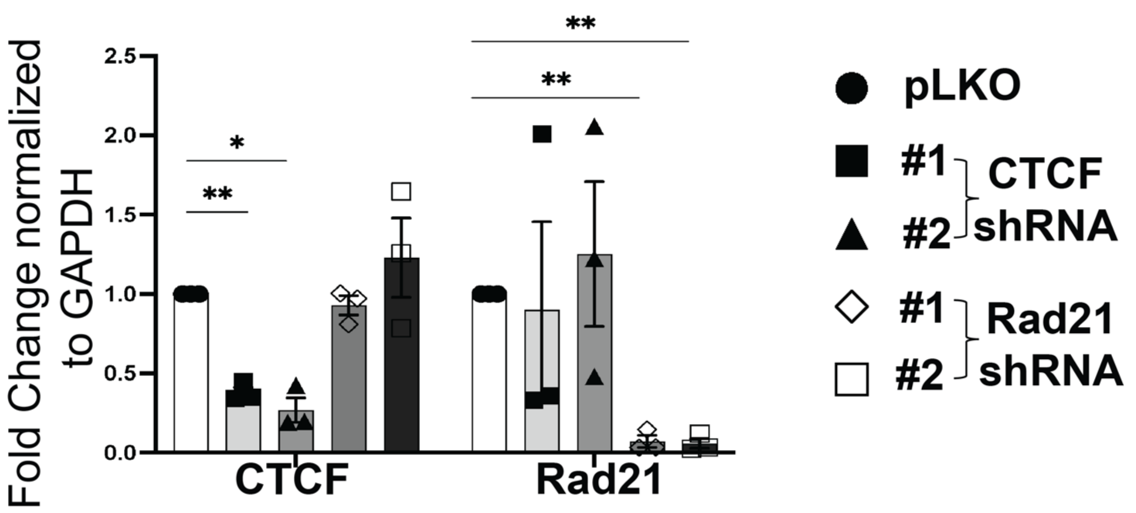


B

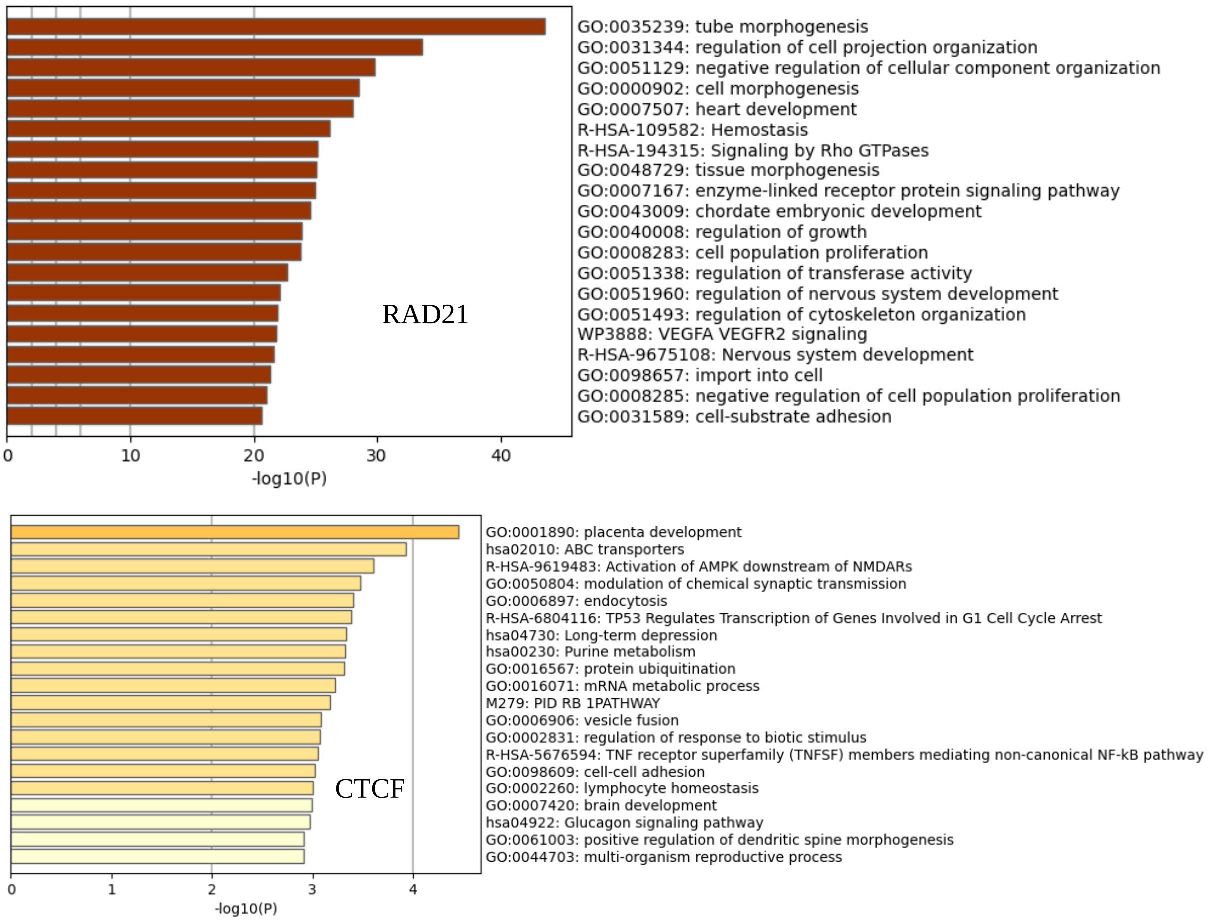


**Figure S11. CTCF and RAD21 regulate gene expression in a chromatin organization-dependent manner. Related to Figure 5.**
(**A**) Knockdown efficiency of CTCF and RAD21 in ECs, as determined by qPCR. N=3 per shRNA; two-sided t tests were used to examine expression difference between groups; *, $p<0.05$.
(**B**) GO/pathway enrichment analysis of DEGs in iECs upon knockdown of Rad21 or CTCF.

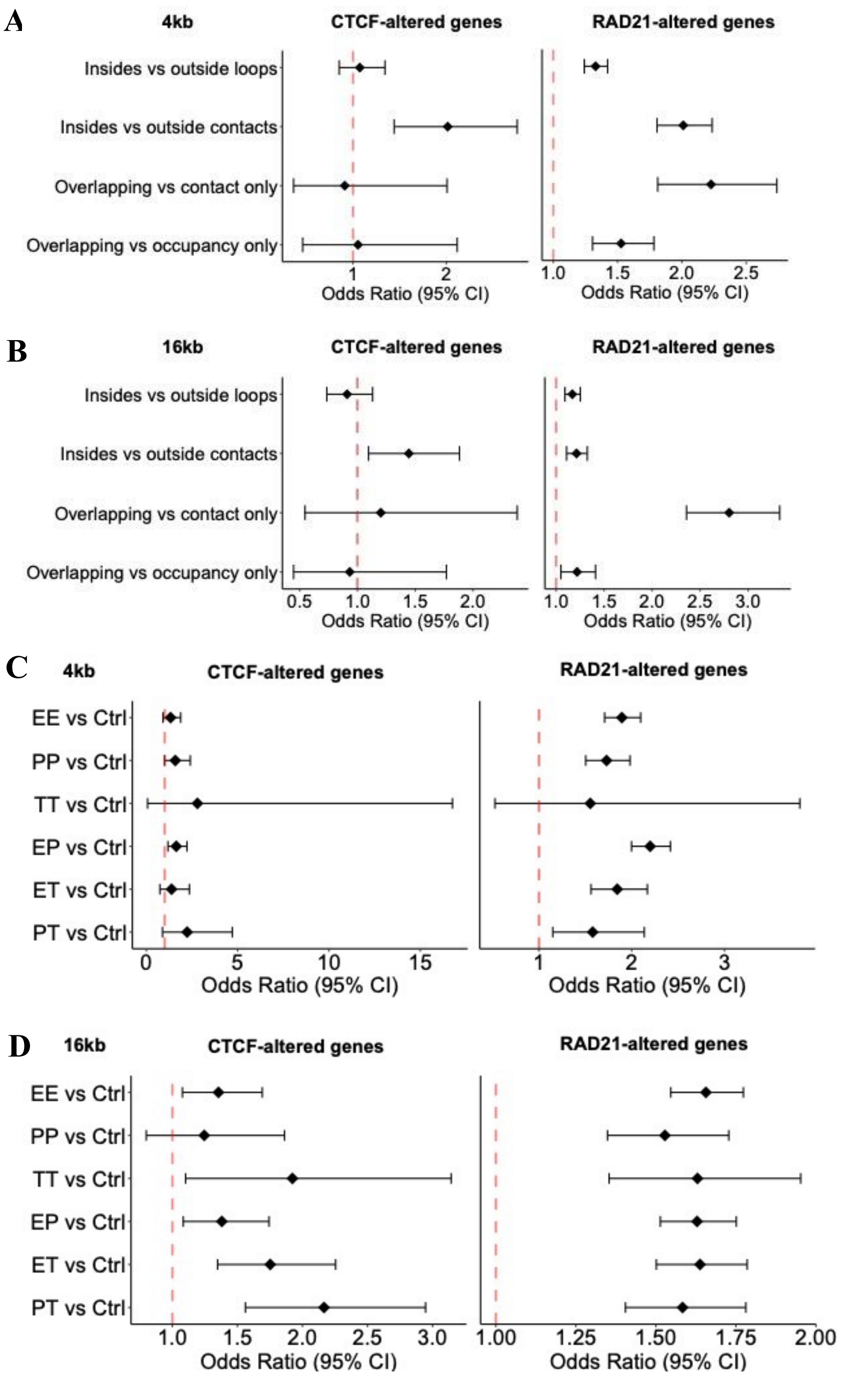


**Figure S12. CTCF and RAD21 regulate gene expression in a chromatin organization-dependent manner. Related to Figure 5, but showing chromatin contacts at 4kb or 16kb resolution instead of the 8kb resolution shown in Figure 5.**

(**A, B**) Genes located in regulatory regions (such as DNA loops, chromatin contact, or near RAD21 or CTCF occupancy sites) are more likely to be altered in iECs, particularly following RAD21 knockdown. Overlapping represent overlapped regions between chromatin contacts and occupancy binding sites of CTCF or RAD21.

(**C, D**) Genes near chromatin contacts involving regulatory interactions are more likely to be altered in response to CTCF or RAD21 knockdown in iECs.

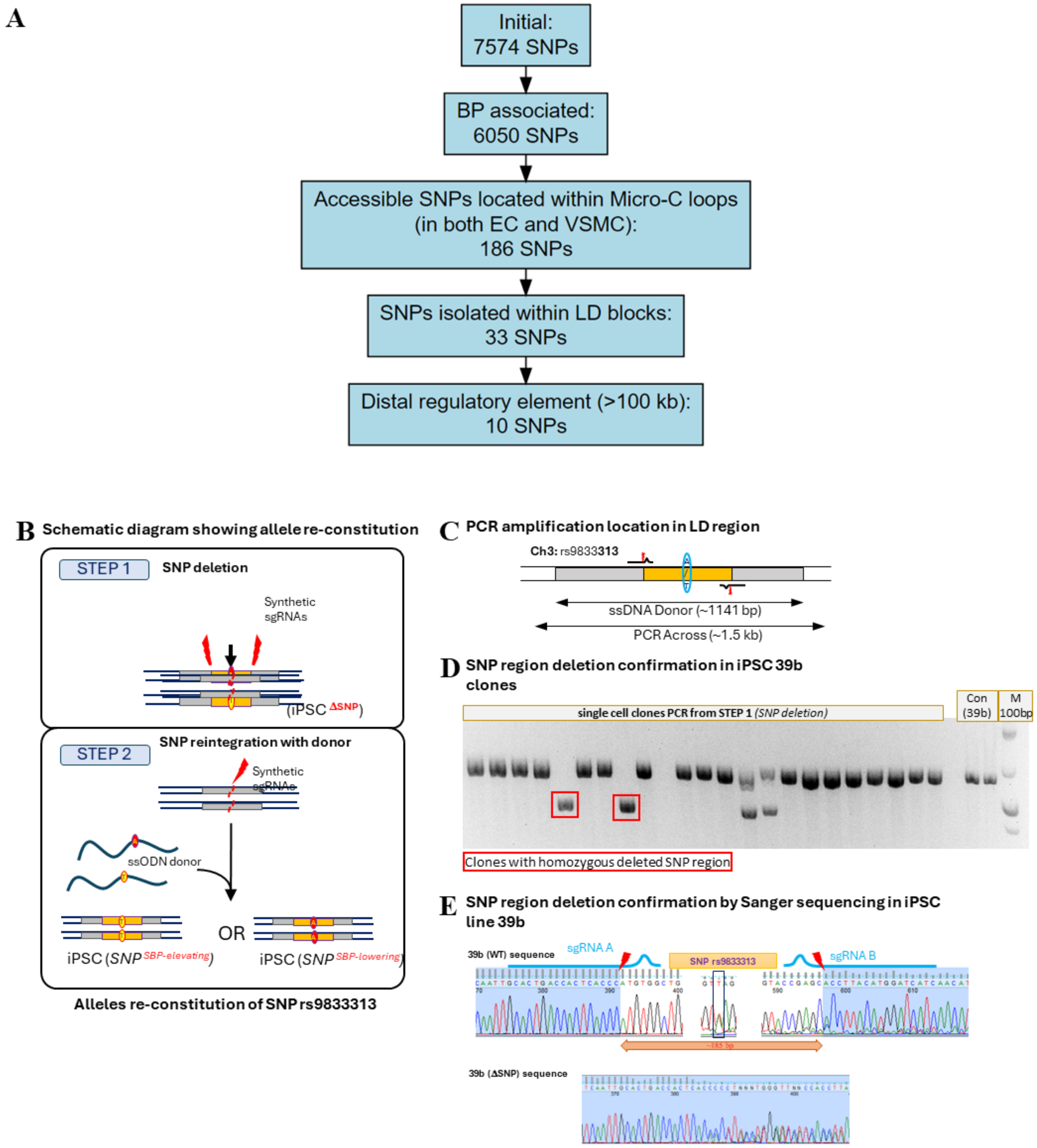


**Figure S13. Two-step genome editing scheme and confirmation of genomic deletion in Step 1. Related to Figure 7.**

(**A**) Pipeline for prioritizing BP-related SNPs for the the proof of principle study.

(**B**) Schematic diagram showing the two-step editing approach (deletion and re-constitution) for generating isogenic hiPSC lines with homozygous allele at SNP rs9833313.

(**C**) Location and size of ssDNA donor and across-region PCR amplicons at rs9833313.

(**D**) Across-region PCR confirmed the deletion of the SNP region, shown as shorter amplicon.

(**E**) rs9833313 region deletion confirmed by Sanger sequencing.

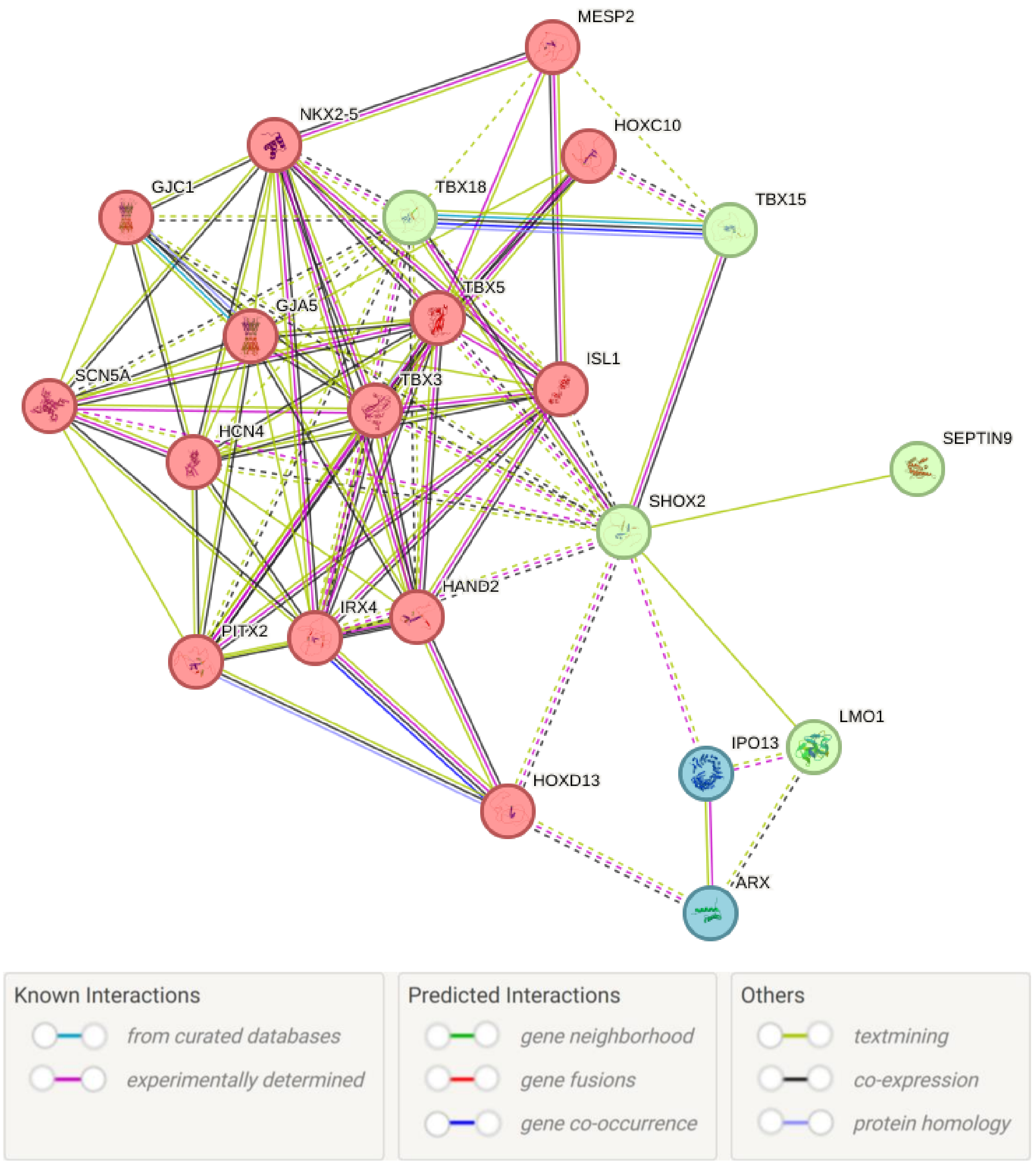


**Figure S14. Genes associated with SHOX2 based on a STRING analysis. Related to Figure 7.** Genes associated with SHOX2 directly or at one step away were shown. Colors of nodes indicate functional clusters. Edges within a cluster are shown as solid lines, and edges between clusters are shown as dotted lines.
